# *Hox*–*Meis*-relayed gene regulatory transition underlies cardiopharyngeal neural crest diversification

**DOI:** 10.1101/2025.11.09.687497

**Authors:** Akiyasu Iwase, Yasunobu Uchijima, Daiki Seya, Mayuko Kida, Hiroki Higashiyama, Kazuhiro Matsui, Akashi Taguchi, Yukihiro Harada, Yunce Wang, Shogo Yamamoto, Shiro Fukuda, Seitaro Nomura, Takahide Kohro, Chisa Shukunami, Haruhiko Akiyama, Masahide Seki, Akinori Kanai, Yutaka Suzuki, Teruhisa Kawamura, Osamu Nakagawa, Hiroto Katoh, Shumpei Ishikawa, Youichiro Wada, Hiroyuki Aburatani, Yukiko Kurihara, Sachiko Miyagawa-Tomita, Hiroki Kurihara

## Abstract

Neural crest cells (NCCs) are multipotent migratory cells essential for cardiac development, yet the lineage trajectories and gene regulatory networks (GRNs) underlying their differentiation in the cardiopharyngeal region remain unclear. Here, we integrate single-cell RNA-seq, spatial transcriptomics, and multiomic analyses to construct a comprehensive map of NCC lineages in developing mouse cardiopharyngeal tissues. We identify a transition from *Hox*-positive pharyngeal NCCs to *Hox*-negative intracardiac populations associated with the outflow tract cushion, accompanied by a shift in *Meis* transcription factor binding and GRN architecture. By contrast, NCCs forming the aorticopulmonary septum and great vessel smooth muscle retain distinct *Hox*-codes. A *Meis2*–*Sox9*–*Scx* GRN defines a skeletogenic progenitor-like intermediate state that gives rise to coronary artery smooth muscle and semilunar valves. Our findings suggest that the loss of *Hox*-dependent regional identity enables pharyngeal NCCs to acquire new fates upon entering the cardiac cushion, providing insight into the developmental origins of coronary and valvular calcification.

## Introduction

Neural crest cells (NCCs) are a multipotent stem cell population that arise from the neural plate border during early vertebrate development^1–3^. As the neural tube forms, NCCs undergo epithelial-mesenchymal transition (EMT), delaminate from its dorsal side, and migrate extensively to differentiate into diverse cell types, including neurons, glia, and melanocytes. NCCs are broadly divided along the anterior-posterior axis into two populations: cranial and trunk NCCs. Cranial NCCs exhibit broader differentiation potential than trunk NCCs and uniquely contribute to mesenchymal (or ectomesenchymal) derivatives such as osteoblasts and chondroblasts, which contribute to craniofacial skeletal structures.

A subset of cranial NCCs also plays a pivotal role in cardiovascular development. Postotic cranial NCCs originating from rhombomeres 6–8, known as cardiac NCCs, migrate through the circumpharyngeal ridge into the caudal pharyngeal arches (PAs) 3, 4, and 6, where they form ectomesenchyme and contribute to the remodeling of the bilaterally symmetrical pharyngeal arch arteries (PAAs) into the asymmetric great arteries, giving rise to their smooth muscle cell (SMC) layers^4,5^. A subpopulation of cardiac NCCs further migrate into the cardiac outflow tract, where they participate in forming the outflow septum and semilunar valves^6^. Ablation of cardiac NCCs in avian embryos results in cardiac malformations, such as persistent truncus arteriosus, underscoring their critical role in heart development^7^. In addition, preotic NCCs from anterior rhombomeres (primarily rhombomere 4) also migrate into the cardiac outflow tract, contributing to coronary artery SMCs and portions of the semilunar valves^8,9^. These findings indicate that a broad range of NCCs participates in cardiac development. Recent advances in single-cell analysis have provided insights into the lineage trajectories and gene regulatory networks (GRNs) governing NCC differentiation. Soldatov et al. demonstrated that, unlike trunk NCCs, cranial NCCs acquire ectomesenchyme potential before delamination from the neural tube depending on the transcription factor (TF) *Twist1*^10^. Similarly, the specification and differentiation of NCCs contributing to cardiovascular development are positionally determined before delamination, and further shaped by environmental cues during migration^11^. Gandhi et al. identified *Tgif1* as a key TF specific to cardiac (postotic) NCCs^12^. However, the mechanisms by which sequential activation of GRNs drives NCC fate diversification in the developing heart in response to spatiotemporal environmental changes remain largely unknown. De Bono et al. elaborated the transition of pharyngeal NCCs through multiple differentiating stages toward SMC fates, identifying *Tbx2* and *Tbx3* as key TFs in this process^13^. They also showed that Tbx1, the gene for 22q11.2 deletion syndrome, regulates pharyngeal NCC differentiation stage transitions non-cell autonomously^13^. However, the relationship between these early NCC derivatives and later-stage intracardiac NCC lineages, as explored by Chen et al.^14^, remains unclear. Furthermore, spatial allocation of each intracardiac NCC derivative is still incomplete.

To bridge these gaps in our understanding of NCC lineages and to elucidate the GRNs driving intracardiac NCC differentiation, we performed single-cell RNA sequencing (scRNA-seq), spatial transcriptomic, and multiomic analyses of NCCs in the pharyngeal and intracardiac region (hereafter referred to as cardiopharyngeal NCCs) using *Wnt1-Cre;R26R-EYFP* mice, a widely used model for NCC lineage tracing^15,16^. By integrating our datasets with publicly available resources, we reconstructed a comprehensive lineage map from early pharyngeal states to diverse cardiac derivatives. We identified a bifurcation in cardiopharyngeal NCC fate: *Hox*-expressing NCCs contribute to great vessel SMCs and the aorticopulmonary septum, while *Hox*-negative NCCs populate the cardiac outflow cushions. This *Hox* status transition was accompanied by a reorganization of GRNs and *Meis* TF binding, leading to the identification of a *Meis2*-, *Sox9*-, and *Scleraxis* (*Scx*)-regulated skeletogenic progenitor-like state that give rise to coronary SMCs and valvular interstitial cells. These findings establish a spatiotemporal framework for cardiopharyngeal NCC lineages and uncover regulatory mechanisms guiding their diversification, with implications for understanding congenital heart defects and pathologic calcification.

## Results

### Cell type heterogeneity within the mouse cardiopharyngeal region

To compare pharyngeal and intracardiac NCCs across developmental stages and characterize their temporal changes, we performed single-cell multiome analysis on pharyngeal and cardiac tissues from E11.5 and E12.5 and cardiac outflow tract tissue from E14.5 and E17.5 *Wnt1-Cre;R26R-EYFP* mouse embryos, in combination with Xenium in situ (E11.5 and E12.5) and Visium spatial transcriptomic (E14.5 and E17.5) analysis on equivalent tissue samples (Figure 1a, b). For single-cell profiling, EYFP-positive NCCs and EYFP-negative non-NCCs were separated using fluorescence-activated cell sorting (FACS) (Figure S1a). Isolated nuclei were subjected to 10x Genomics RNA+ Assay of Transposase Accessible Chromatin (ATAC) Multiome analysis (Figure 1b). After quality control filtering, we obtained scRNA-seq and single-cell ATAC sequencing (scATAC-seq) data from 4,876 NCC nuclei and 4,544 non-NCC nuclei.

**Figure 1.**
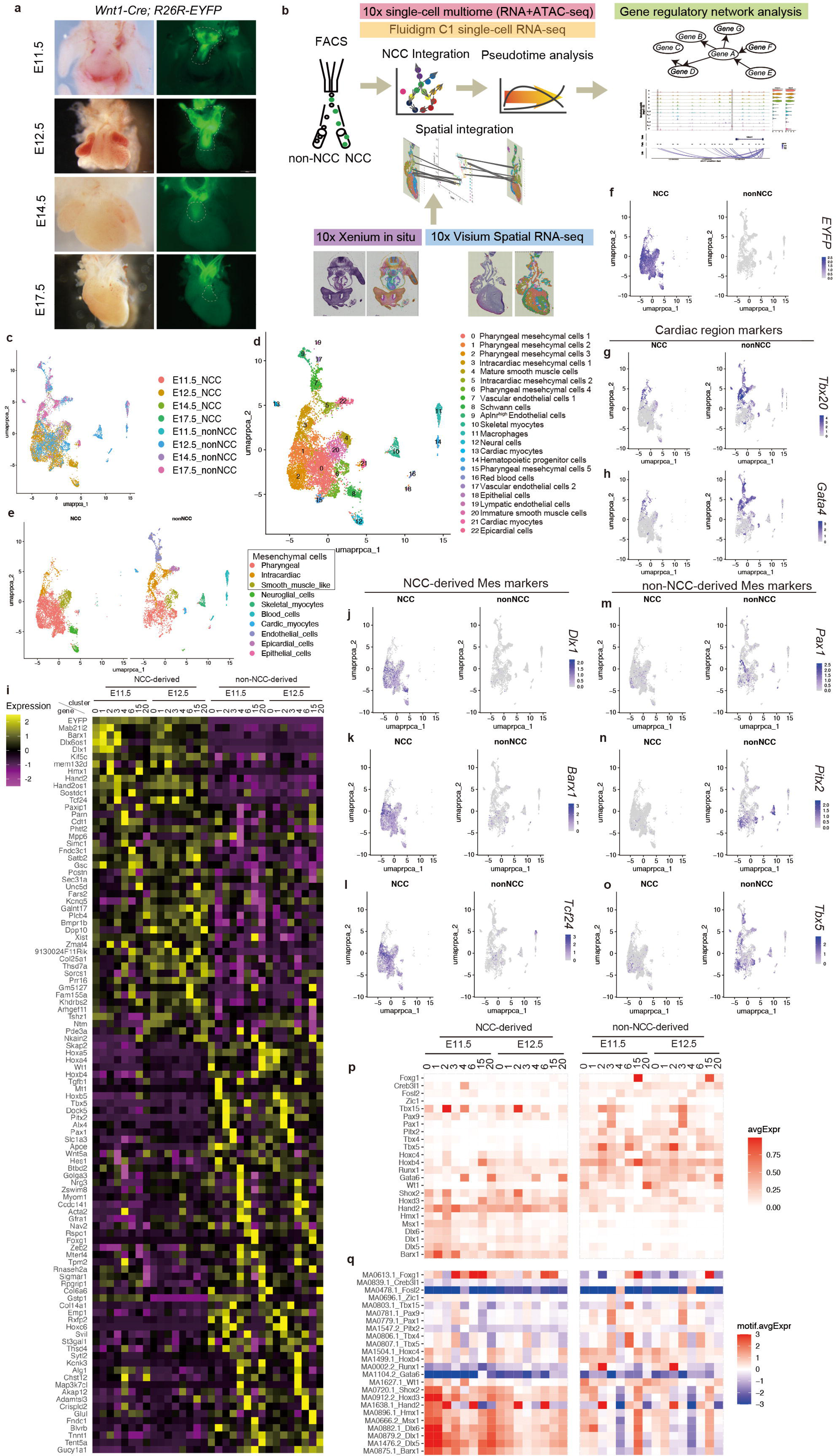
Heterogeneity of NCC- and non-NCC-derived cell populations during cardiopharyngeal development. (a) Localization of NCCs labeled with *Wnt1-Cre;R26R-EYFP* in cardiopharyngeal tissues. Regions outlined by dashed lines were dissected for single-cell analyses. (b) Workflow of single-cell and spatial transcriptomic analysis, with a focus on NCCs. (c-e) UMAP plots of scRNA-seq data, colored by embryonic stage and cell type (c), fine cluster identity (d), and coarse cluster identity (e). (f-h) Feature plots showing expression of EYFP (f) and cardiac markers *Tbx20* (g) and *Gata4* (h). (i) Heatmap of differentially expressed genes (DEGs) between NCC-derived and non-NCC-derived mesenchymal cells. For comparison between the two populations at E11.5 and E12.5, DEGs with an average log2 expression > 0.15 were ranked by adjusted p-value from the Wilcoxon rank-sum test with Bonferroni correction, and the top 5 genes were selected for each cluster (0, 1, 2, 3, 4, 6, 15, and 20 in panel d). Expression values were averaged across integrated clusters. (j-o) Feature plots showing expression of NCC-derived mesenchymal markers *Dlx1* (j), *Barx1* (k), and *Tcf24* (l), and non-NCC-derived mesenchymal markers *Pax1* (m), *Pitx2* (n), and *Tbx5* (o). (p-q) Heatmaps comparing differentially expressed TF genes (p) and the enrichment of their binding motifs in open chromatin regions (q) between NCC-derived and non-NCC-derived mesenchymal cells at E11.5 and E12.5.

Unsupervised clustering with the uniform manifold approximation and projection (UMAP) separated a total of 9,420 cells into 23 distinct clusters based on their transcriptomic profiles and TF motif enrichment (Figure 1c-e, Figure S1b-d, and Table S 1, 2). The majority of NCCs formed a broad mesenchymal population interspersed with non-NCCs (Figure 1c-e), while a subset of NCCs segregated into two discrete clusters corresponding to glial and neural cell populations (Figure 1c-e and Figure S1b-d). Only a few NCCs were detected within cardiomyocyte clusters, which were predominantly composed of non-NCCs, consistent with previous reports demonstrating NCC differentiation into cardiomyocytes^15,16^. The overall proportion of cardiomyocytes was low, likely reflecting the restricted sampling of the cardiac outflow tract region (Figure 1a). The remaining clusters were composed almost exclusively of non-NCCs (Figure 1e, f).

Within the mesenchymal compartment, Clusters 4 and 20 constituted a smooth muscle-like population as indicated by high expression of smooth muscle markers such as *Myh11*, *Cnn1*, and *Acta2* (Figure S1b). Cluster 3 and 5 were distinguished by enrichment of cardiac markers such as *Tbx20* and *Gata4* (Figure 1g, h), consistent with an intracardiac mesenchymal population distinct from other pharyngeal mesenchymal populations (Figure 1d, e).

Among mesenchymal populations at E11.5 and E12.5, NCC-derived and non-NCC-derived cells were distinguished by a set of genes differentially expressed (Figure 1i). NCC-derived cells (ectomesenchyme) exhibited high expression of TF genes such as *Dlx1, Barx1, Mab21l2, Satb2*, and *Tcf24* (Figure 1j-l, o and Table S3). In contrast, non-NCC-derived mesenchymal cells were characterized by the expression of *Pax1, Tbx5,* and *Pitx2* (Figure 1n-o and Table S3). Consensus binding motifs of these TFs were correspondingly enriched in the open chromatin regions of their respective cell populations (Figure 1p, q and Table S4). These distinctions were less pronounced in comparisons between NCCs and non-NCCs within the cardiac outflow tract at E14.5 and E17.5 (Figure S1e, f and Table S3, 4).

### Spatial allocation of NCC clusters through the integration of single-cell and spatial transcriptomic analysis

To spatially resolve NCC-derived cell types, we applied Xenium in situ hybridization to transverse sections of the cardiopharyngeal region at E11.5 and E12.5, using customized panel probes. After assigning a total of 1,003,927 cells to 39 cell types with histological validation by hematoxylin-eosin staining (Figure 2a-e, Figure S2a-j, and Table S5), the spatially annotated datasets were then integrated with single-cell multiome data using Tangram, a deep learning method aligning single-cell data to spatial data^17^, allowing the estimation of gene expression beyond the probe set. This analysis estimated EYFP-expressing NCCs to be distributed within the pharyngeal region around the great arteries and the trachea, the cardiac outflow tract endocardial cushion, and peripheral nerve tissues (Figure 2f, g). These spatial distributions were consistent with lacZ-expressing cells in *Wnt1-Cre;R26R-lacZ* embryos (Ref. ^18^ and Figure S2k, l), validating the observed NCC-derived cell populations. Furthermore, these regions were enriched for *Tcf24*, one of the genes identified as NCC-specific (Figure 2h, i).

**Figure 2.**
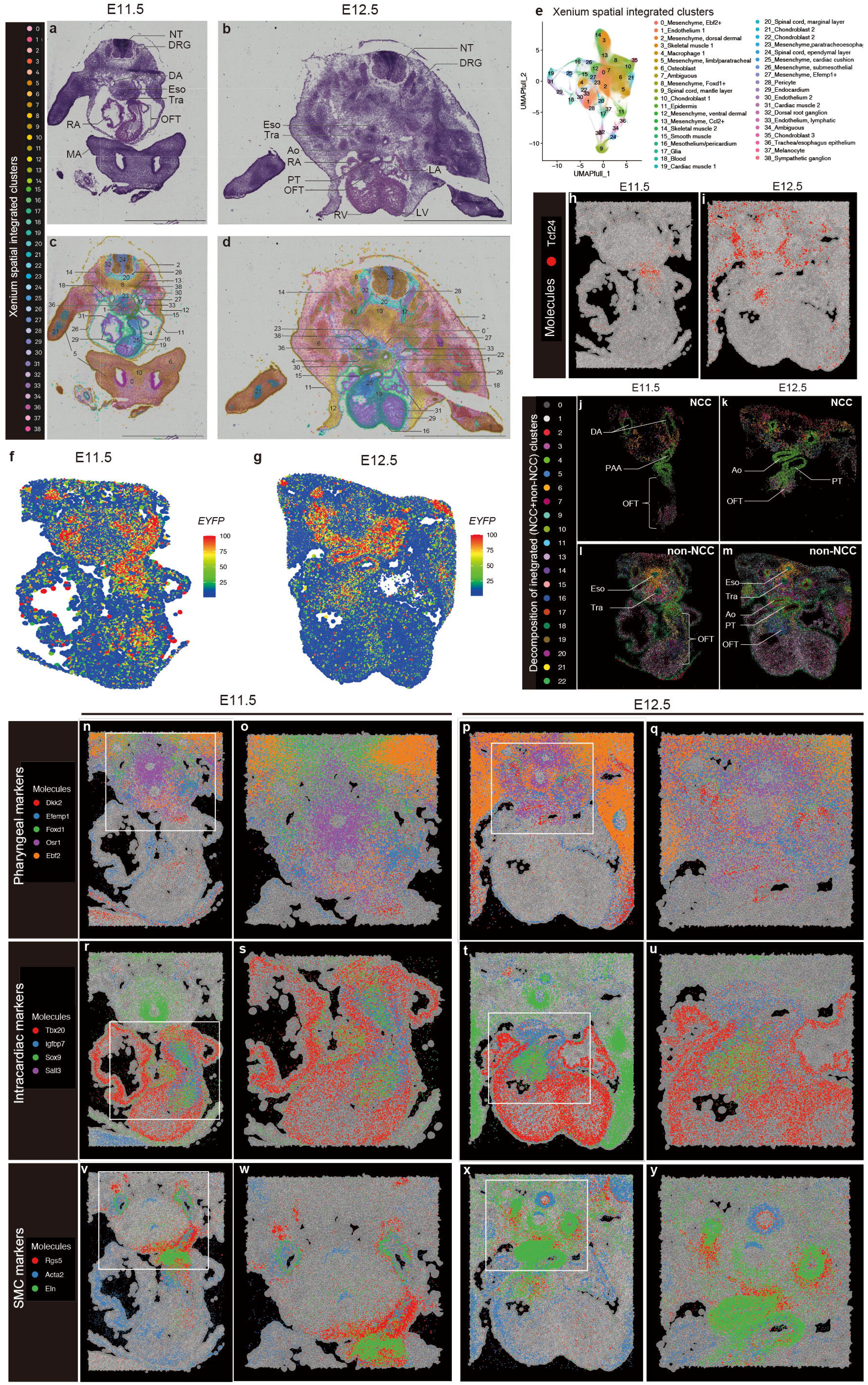
Spatial distribution of NCCs with distinct transcriptomic signatures. (a-d) Xenium datasets from cardiopharyngeal tissues at E11.5 (a, c) and E12.5 (b, d) with hematoxylin-eosin staining. Ao, aorta; DA; dorsal aorta; DRG, dorsal root ganglion; Eso, esophagus; LA, left atrium; LV, left ventricle; MA, maxillary arch; NT, neural tube; OFT, outflow tract; PT, pulmonary trunk; RA, right atrium; RV, right ventricle; Tra, trachea. Scale bars, 2mm. (e) UMAP plots of 1,003,927 cells profiled by Xenium analysis, colored by Xenium spatial integrated clusters based on gene expression signatures. (f, g) Tangram-based estimation of *EYFP* expression in cardiopharyngeal tissues at E11.5 (f) and E12.5 (g). (h, i) Xenium images showing *Tcf24* expression in cardiopharyngeal tissues at E11.5 (h) and E12.5 (i). (j-m) RCTD decomposition of the scRNA-seq dataset (Figure 1d) into NCC (j, k) and non-NCC (l, m) components, spatially mapped onto E11.5 (j, l) and E12.5 (k, m) Xenium datasets. (n-y) Xenium images showing regional and cell-type markers that delineate NCCs in the cardiopharyngeal region at E11.5 (n, o, r, s, v, w) and E12.5 (p, q, t, u, x, y). NCCs are classified into pharyngeal mesenchymal cells (n–q), intracardiac mesenchymal cells (r–u), and SMCs (v–y). Boxed regions in n, p, r, t, v, and x are magnified in panels o, q, s, u, w, and y, respectively. Note that these markers are not exclusive to NCC derivatives.

To refine spatial localization further, we decomposed the scRNA-seq dataset into NCC and non-NCC components and mapped them onto the Xenium dataset using the robust cell type decomposition (RCTD) method^19^ (Figure 2j-m). This approach categorized NCCs in the E11.5–E12.5 cardiopharyngeal region into three major populations based on regional identity and marker gene expression: (1) pharyngeal mesenchymal cells (*Osr1*^high^ and *Ebf2*^high^) (Figure 2n-q), (2) intracardiac mesenchymal cells (*Sox9*^high^ and *Tbx20*^high^) (Figure 2r-u), and (3) SMCs (*Acta2*^high^ and *Eln*^high^) (Figure 2v-y). These groups corresponded approximately to the mesenchymal NCC clusters identified in the single-cell transcriptomic analysis, and were further validated by the expression of well-established cell type-specific markers (Figure S1a and Table S1). Notably, *Tbx20* expression was broadly detected in cardiac tissues beyond NCC-derivatives and was distinctly demarcated from the pharyngeal regions at the pericardial reflection (Figure 2r-u).

### Comprehensive mapping of the landscape of cardiopharyngeal NCC lineages

These distinct regional expression patterns observed lead us to investigate how successive changes in gene expression drive NCC differentiation into diverse lineages. To achieve a comprehensive understanding of cardiopharyngeal NCC lineages, we integrated our data with the publicly available scRNA-seq datasets of NCCs from E8.5 embryos to postnatal day 7 (P7) hearts^13,14^. Two-dimensional UMAP separated a total of 67,208 cells into 28 distinct clusters (Cs) based on their transcriptomic profiles (Figure 3a, b, Figure S3a, and Table S6).

**Figure 3.**
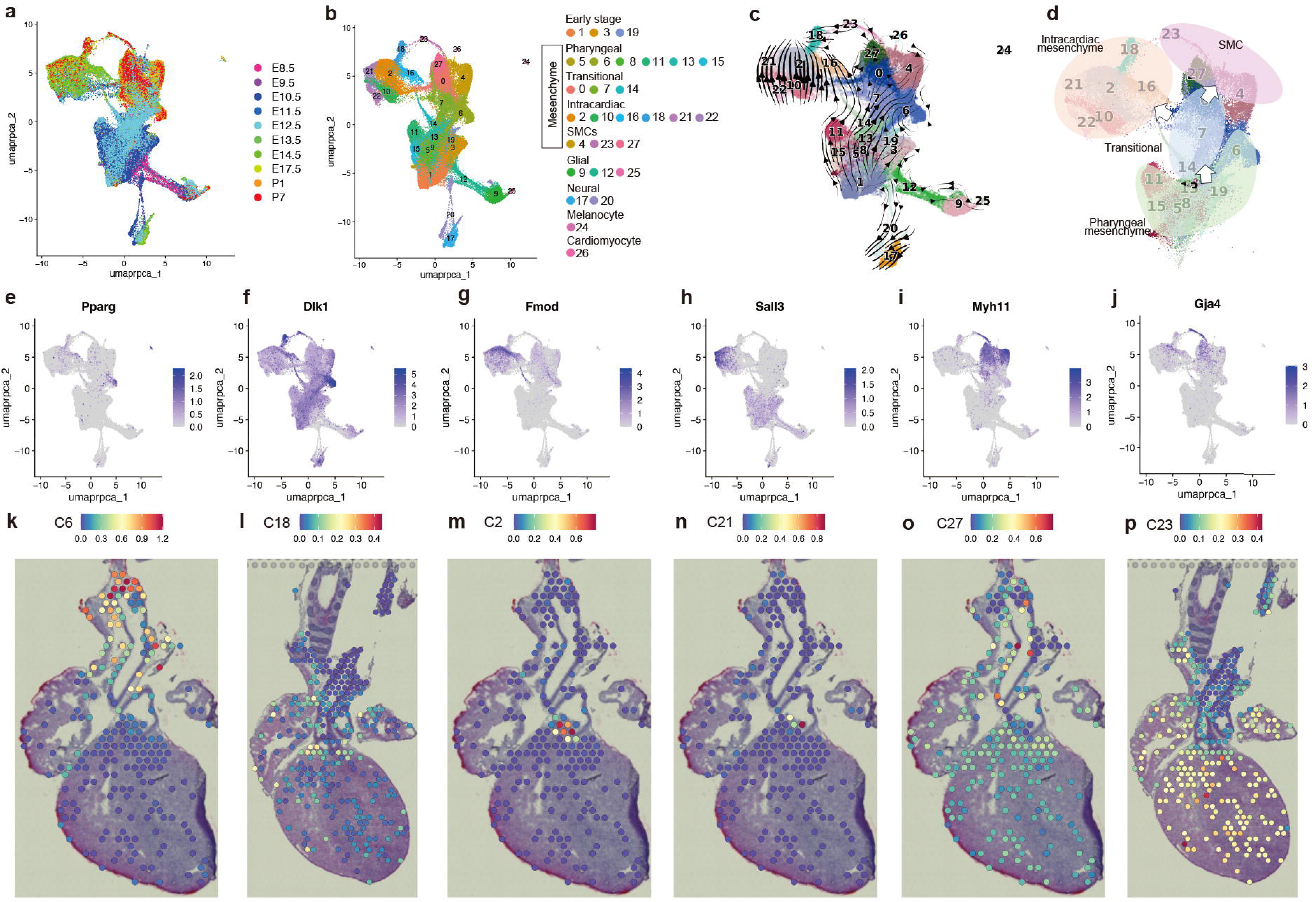
Comprehensive reconstruction and spatial characterization of NCC-derived cardiopharyngeal mesenchymal lineages using integrated scRNA-seq datasets. (a, b) UMAP plots of integrated scRNA-seq data from craniopharyngeal NCCs, colored by embryonic stage (a), and cluster identity (b). (c) RNA velocity flow embedded on the integrated UMAP. (d) Categorization and inferred lineage relationships among NCC-derived mesenchymal clusters. (e-j) Feature plots showing representative marker gene expression for mesenchymal clusters; *Pparg* (e), *Dlk1* (f), *Fmod* (g), *Sall3* (h), *Myh11* (i), and *Gja4* (j). (k-p) Spatial localization of representative clusters on the Visium datasets; C6 (k), C18 (l), C2 (m), C21 (n), C27 (o), and C23 (p).

NCC-derived lineages included glial (C9, C12, C25), neural (C17, C20), melanocyte (C24), and cardiomyocyte (C26) clusters, as well as early-stage clusters containing the neural tube (C1, C3, C19) (Figure 3b, Figure S3a-g, and Table S6). The remaining 18 mesenchymal clusters were broadly categorized into four major groups: pharyngeal mesenchyme (C5, C6, C8, C11, C13, C15), intracardiac mesenchyme (C2, C10, C16, C18, C21, C22), SMCs (C4, C23, C27), and transitional states (C0, C7, C14), corresponding to the categories described above (Figure S3b, e-k, and Table S6). RNA velocity analysis in conjunction with developmental context, inferred global lineage relationships among these groups (Figure 3c, d), consistent with developmental trajectories in vivo.

The pharyngeal mesenchyme group (*Osr1*^high^ and *Ebf2*^high^) was mainly composed of ectomesenchymal cells at early stages (Figure S3h). Within this group, C6 showed high expression of adipocyte differentiation markers *Pparg*, *Cebpa*, and *Mfap5* (Figure 3e and Table S6), indicating that this cluster represents a population containing progenitors of periaortic brown adipocytes^20^. The remaining clusters displayed considerable overlap and retained features of immature mesenchyme.

The SMC clusters, which were continuous with the pharyngeal mesenchyme via transitional populations in the UMAP, were identified by high expression of the mature SMC marker *Myh11* (Figure 3i). Differential gene expression analysis further distinguished individual clusters (Figure S4a-g). Among these, C27 displayed a transcriptomic profile characteristic of the great artery SMCs, including high expression of *Sost* (Figure S3j). C4 was enriched for *Tfap2b* and *Ptger4* (Figure S3j), markers of the ductus arteriosus SMCs^21,22^, supporting its annotation. C0 and C7 likely represent transitional states between pharyngeal mesenchyme and differentiated lineages, potentially bifurcating toward great artery SMCs or cardiac cushion mesenchyme (Figure S3a and Table S6). C23 was characterized by high expression of *Gja4*, a marker of coronary artery SMCs, along with pericyte markers *Kcnj8* and *Rgs5* (Figure 3j and Figure S3k), corresponding to the cluster similarly annotated by Chen et al^14^. In addition, C23 was also distinguished from C4 and C27 by its expression of *Reln* (Figure S4d).

Immunostaining supported these cluster annotations. Sost expression is observed in great artery SMCs but not in ductus arteriosus and coronary artery SMCs, whereas Myh11 expression was higher in ductus arteriosus and coronary artery SMCs than in aortic SMCs (Figure S4f-m). Furthermore, Reln expression was restricted to coronary artery SMCs (Figure S4n-s).

Notably, the UMAP revealed a continuum between C23 and C18 within the intracardiac mesenchyme population. Given previous findings that the proximal coronary artery SMCs originate from preotic NCCs^8^ and that pericytes give rise to coronary artery SMCs^23^, this connection likely represent a differentiation trajectory from intracardiac mesenchyme to coronary artery SMCs via a pericyte-like intermediate stage.

Within the intracardiac mesenchyme group, C18 exhibited high *Dlk1* and *Tcf21* expression (Figure 3f and Figure S3i). Immunostaining for Dlk1 confirmed that this population represents mesenchymal cells localized to the cushion-derived subvalvular region (Figure S3l).

C16 was distinguished by high expression of *Penk* and *Sfrp2* (Figure S3i and Table S6), corresponding to the cluster annotated as the aorticopulmonary septum in the previous study by Chen et al^14^. This cluster also exhibited robust expression of mesenchymal markers, including *Postn,* similar to other intracardiac clusters (Figure S3i). In addition, C16 showed relatively high expression of *Tcf24* and low expression of *Vegfc* compared with the other intracardiac clusters (Figure S5a-e). These gene expression features of the aorticopulmonary septum were further validated by Xenium in situ hybridization (Figure S5f-j).

Unlike other intracardiac NCCs that populate the distal outflow tract cushions, the aorticopulmonary septum originates as a protrusion from the dorsal wall of the aortic sac and is primarily derived from NCCs residing in PA4 and PA6^6,24–26^. This septal structure subsequently fuses with the distal outflow tract cushions to partition the common arterial trunk into the aortic and pulmonary channels. Consistent with this developmental origin, C16 was enriched for the expression of *Hox4* and *Hox5* paralogs (Figure 3t, u), indicating that NCCs in this population retain their *Hox* code, in contrast to other intracardiac NCCs, in which most *Hox* genes were downregulated (see later details).

The remaining clusters C2, C10, C21, and C22 showed substantial overlap and likely represent valve-forming mesenchyme, as suggested by their high expression of cushion and valve interstitial markers such as *Hapln1* and *Postn* (Figure S3i). Among these, C2 and C21 were further distinguished by enriched expression of *Fmod* and *Sall3*, respectively (Figure 3g, h).

To further investigate the spatial distribution and interconnectivity of intracardiac and related cell clusters, we mapped them onto Visium spatial transcriptomic datasets from E14.5 and E17.5 mouse hearts (Figure S6a-l and Table S7) using RCTD in full mode, enabling deconvolution of each spatial spot into multiple cell types (Figure 3k-p and Figure S7). This analysis largely validated the assigned spatial identities. C6 was localized adjacent to the aorta (Figure 3k), in line with its role as a source of NCC-derived periaortic brown adipocytes^20^. Within the intracardiac mesenchyme, C18 and C2 were enriched in the subvalvular region (Figure 3l, m), while C21 was concentrated in the semilunar valves (Figure 3n). C10 and C22 were broadly distributed across valvular and subvalvular regions (Figure S7). Given their enrichment in the S or G2/M phases at earlier developmental stages (Figure S3d), they likely represent immature, proliferating mesenchymal cells. Based on developmental timing and connectivity in the UMAP, we infer that intracardiac NCCs within C10 and C22 differentiate via C2 to valvular (C21) and subvalvular (C18) interstitial cells, the latter serving as progenitors of coronary artery SMCs (C23) (Figure 3c, d). C27 and C23 were localized to the ascending aorta and the cardiac base, respectively (Figure 3o, p), consistent with its identity as a pericyte as well as coronary artery SMC population.

### *Hox* code patterns distinguish NCC subpopulations during cardiopharyngeal development

NCCs in the PAs and associated arteries, excluding the first PA (PA1), exhibit distinct, nested patterns of *Hox* gene expression along the anterior-posterior axis^27,28^. Based on this regional identity, we decoded the *Hox* code pattens of cardiopharyngeal NCC populations using integrated scRNA-seq data (Figure 4a-e and Figure S8). Although RNA-seq may underestimate *Hox* expression due to potential false negatives, our analysis effectively captured key regional signatures, as exemplified by ductus arteriosus SMCs (C4), exhibiting enrichment for *Hox4* and *Hox5* paralogs (Figure 4i, j). These *Hox*-code populations were similarly distributed among pharyngeal and transitional NCCs, great artery SMCs, and aorticopulmonary septum NCCs (Figure 4k and Figure S8). By contrast, most intracardiac NCCs, including those giving rise to coronary artery SMCs, were largely *Hox*-negative with the exception of the aorticopulmonary septum (Figure 4k and Figure S8). Comparative analysis of *Hox* paralog expression across NCC subtypes further supported this classification (Figure 4i, j and Figure S8).

**Figure 4.**
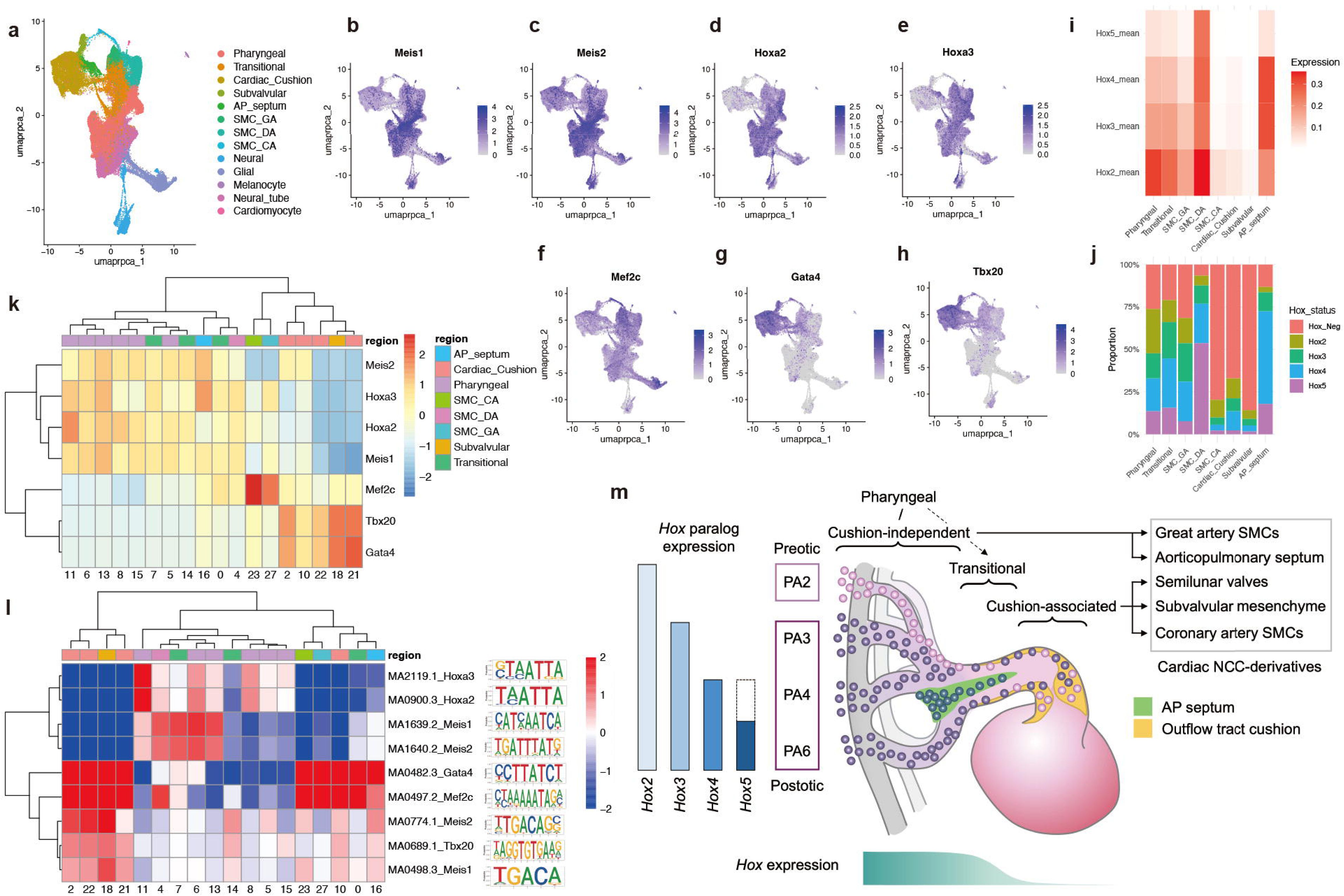
TF expression and motif enrichment of NCC-derived cardiopharyngeal mesenchymal lineages. (a) Subcategorization of NCC clusters reflecting heterogeneity within intracardiac and SMC populations. (b-h) Feature plots showing expression of *Meis1* (b), Meis2 (c), *Hoxa2* (d), *Hoxa3* (e), *Mef2c* (f), *Gata4* (g), and *Tbx20* (h). (i) Heatmap displaying anterior *Hox* gene expression across NCC-derived mesenchymal subtypes defined in (a). (j) Classification of mesenchymal subtypes based on *Hox* codes: Hox-negative; Hox2 (expressing only *Hox2* paralogs); Hox3 (expressing *Hox3* paralogs without *Hox4/5*); Hox4 (expressing *Hox4* paralogs without *Hox5*); and Hox5 (expressing *Hox5* paralogs, with or without anterior *Hox* genes). (k-l) Heatmaps showing (k) TF gene expression and (l) enrichment of corresponding DNA-binding motifs in open chromatin regions across mesenchymal subtypes. (m) Schematic summarizing NCC lineage relationships, spatial contributions, and corresponding *Hox* expression dynamics during cardiopharyngeal development.

Given that NCCs populating the cardiac outflow cushion originate from preotic and postotic rhombomeres with substantial *Hox* codes^8^, the observed downregulation of *Hox* genes likely occurs upon migration into the outflow cushion. This is further substantiated by the reduced enrichment of Hox-binding motifs in open chromatin regions of cushion-associated NCCs compared to pharyngeal and transitional NCCs (Figure 4l).

Collectively, these findings delineate two categories of NCCs contributing to the cardiovascular development: (1) cushion-associated NCCs, which give rise to the semilunar valves, subvalvular mesenchyme, and coronary artery SMCs, characterized by *Hox* gene downregulation, and (2) cushion-independent NCCs, which contribute to the aorticopulmonary septum and great artery SMCs, and maintain regional *Hox* gene expression (Figure 4m).

To further delineate Hox code patterns associated with pharyngeal arch origin, we stratified the integrated UMAP by distinct *Hox* expression profiles (Figure S9a). Cells expressing any *Hox2* paralog, but lacking *Hox3–5* paralogs, were defined as PA2-derived preotic NCCs, whereas cells expressing any of *Hox3–5* paralogs were classified as PA3/4/6-derived postotic NCCs. Preotic, postotic, and *Hox*-negative populations were then projected onto the integrated UMAP across developmental stages (E10.5–E14.5). Trajectory inference indicated that transitions toward intracardiac mesenchyme occur earlier in preotic cells (E10.5) than in postotic cells (E11.5), consistent with their known sequential migration into the cardiac cushion^8^. From E12.5 to E14.5, postotic cells showed a progressive emergence of the aorticopulmonary septum–associated cluster C16 from transitional states. Notably, the proportion of *Hox*-negative cells increased within intracardiac mesenchyme, except in C16 where Hox expression was retained, supporting the notion that *Hox* genes are broadly downregulated in cushion-associated intracardiac NCCs (Figure 4k, S9).

Lineage trajectories were further investigated using Slingshot^29^ and CellRank2^30^-integrated RNA velocity analysis. At E10.5–E11.5, the transitional continuum (C14, C7, and C0) diverged into intracardiac cushion mesenchyme (C2), great artery SMCs (C27), and ductus arteriosus SMCs (C4) (Figure S9b,c). Along this diversification, TFs characteristic of transitional states, such as *Foxf1* in C14 and C7, were replaced by lineage-specific TFs, including *Gata4* and *Tbx20* in C2 (Figure S9d). After E12.5, a trajectory toward the aorticopulmonary septum–associated cluster C16 bifurcated from C14, which directed to great artery SMCs (C27) and ductus arteriosus SMCs (C4) through C7 and C0 (Figure S9e,f). C16 shared expression of *Gata4* and *Tbx20* with C2 (Figure S9g), but retained *Hox* gene expression (Figure 4k, S8). The great artery SMC lineage was marked by *Sost* expression (Figure S9g), consistent with immunostaining results (Figure S4i,l). The ductus arteriosus SMC trajectory was characterized by induction of *Tfap2b* and reactivation of *Foxf1* (Figure S9g).

### Partner switching of Meis proteins reflects functional transition in cardiopharyngeal NCCs

Hox proteins function through interactions with TALE (three-amino-acid loop extension) homeodomain proteins, including Meis and Pbx^31,32^. Of the two primary Meis binding motifs, the TGATT(T/G)AT octamer and the TGACAG hexamer, Hox-Pbx-Meis complexes preferentially bind to the octameric motif, whereas Meis proteins interacting with non-Hox TFs tend to bind to the hexameric motif^33^.

In line with the observed downregulation of *Hox* gene expression and diminished accessibility of Hox-binding motifs in cushion-associated NCCs, enrichment of Meis-binding octameric motifs (MA1639.2 and MA1640.2) was significantly decreased (Figure 4l). Conversely, enrichment of the hexameric motifs (MA0498.3 and MA0774.1) was significantly increased compared to pharyngeal and transitional NCCs (Figure 4l). Expression levels of *Meis1* and *Meis2* remained unchanged (Figure 4k), suggesting that the observed shift in motif enrichment reflects changes in transcriptional partners rather than Meis protein abundance.

Accompanying this switch, both expression and binding motif accessibility of *Gata4* and *Tbx20*, non-Hox TFs critical for cardiac development, were upregulated in cushion NCCs (Figure 4k, l). These results support a model in which Meis proteins undergo a functional transition during cardiac cushion-associated NCC differentiation, shifting from Hox-dependent to Hox-independent transcriptional regulation via alternative partner interactions and DNA binding preferences.

### Inference of GRNs driving cardiac NCC differentiation

Dynamic reorganization of GRNs, including the downregulation of *Hox* gene expression and a switch in Meis binding partners from *Hox* to non-*Hox* TFs, likely underlies distinct differentiation programs of NCCs during cardiac development. To elucidate these GRNs, we utilized additional scRNA-seq datasets from EYFP^+^ NCCs isolated at E11.5, E12.5, E14.5 and E17.5 from *Wnt1-Cre;R26R-EYFP* mouse hearts, generated using the Fluidigm C1 platform. Although these datasets contained fewer cells, their high read depth made them suitable for GRN inference.

We projected the Fluidigm C1 onto the integrated UMAP, categorizing cells into cushion-associated (C2, C10, C18, C21, C22, C23), cushion-independent (C0, C4, C16, C27), and pharyngeal or transitional (C6, C7, C14) populations (Figure 4a and Figure S10a). To infer GRNs, we applied a nonparametric Bayesian network approach using the NNSR algorithm^34^. Initial analysis via SCENIC^35^, focused on TF binding sites within ±10kb of transcription start sites (TSSs), identified 269,646 potential TF–target gene links, of which 18,655 links surpassed the frequency threshold. Filtering for genes annotated as TFs yielded a network of 560 nodes and 1,588 edges (Table S8). Using the Linkcomm algorithm^36^, we partitioned this network into 109 overlapping communities (Table S9). Subnetworks connecting TFs within each community to their first-edge target genes were defined as “overall communities” (OCs). OC enrichment scores were then computed for each cell (scRNA-seq) or Visium spot (Figure 5a-c and Figure S10b). Then, we performed random forest classifier to extract key OCs involved in regulating lineage commitment (Figure S10c, d). Mapping OC dynamics onto the integrated UMAP and lineage annotations revealed distinct GRN architectures among NCC subpopulations (Figure 5c). Notably, cushion-associated and cushion-independent lineages exhibited divergent OC profiles, reflecting distinct transcriptional programs, even when both contribute to SMC differentiation (Figure 5d).

**Figure 5.**
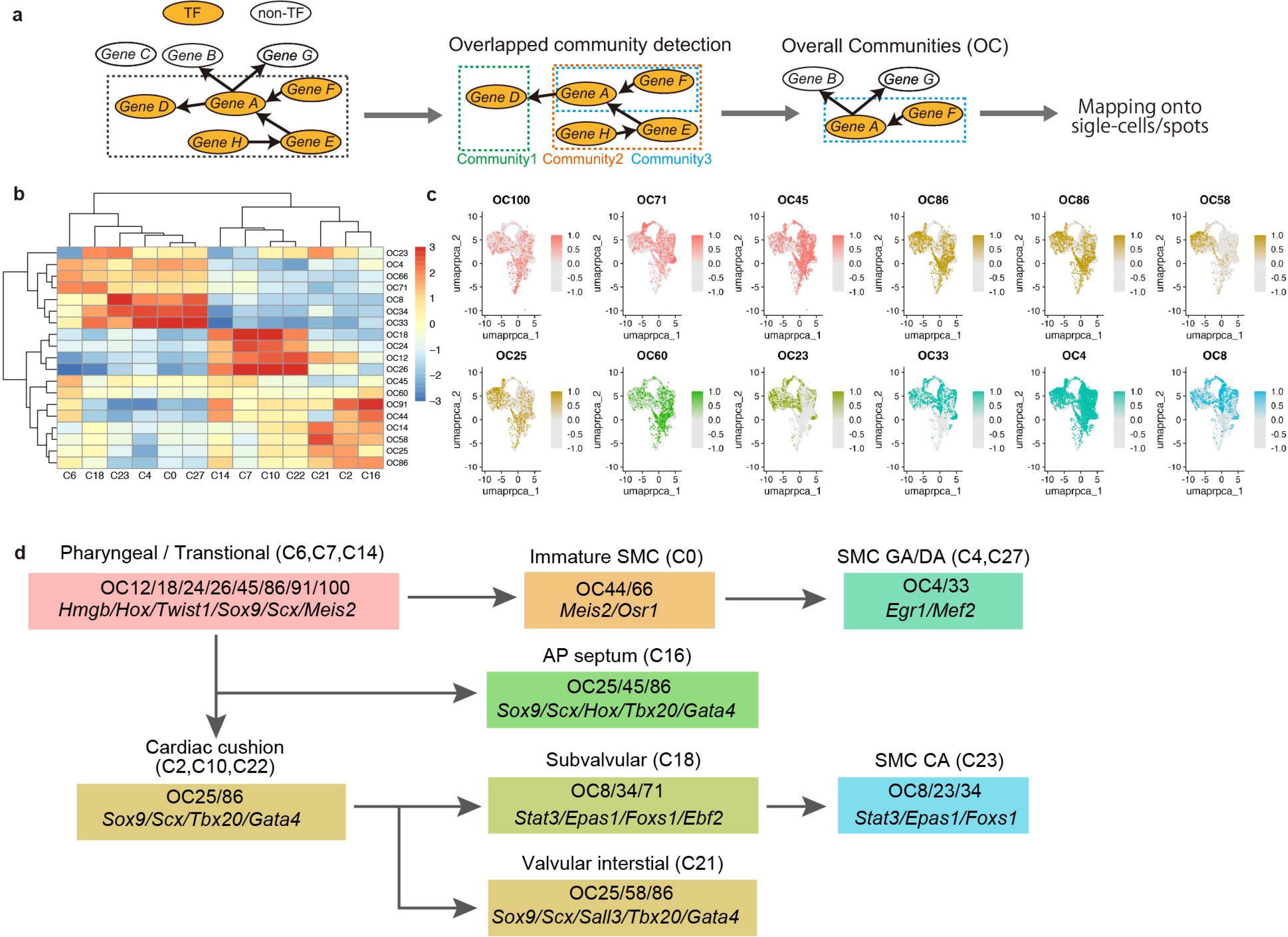
GRNs involved in cardiopharyngeal NCC fate determination. (a) Scheme of GRN analysis using Fluidigm C1 scRNA-seq data. Dashed-line boxes indicate the TF subnetwork, which was divided into overlapping communities. Each TF is connected to its first-edge targets to define “overall communities” (OCs). (b) Heatmap showing the enrichment of representative OCs across integrated clusters. Color scale represents z-scored enrichment values. (c) UMAP plots showing representative OC enrichment patterns, colored by cluster identity as in (d) with intensity indicating enrichment levels. (d) Schematic representation of NCC lineage trajectories annotated with OCs and key TFs.

In pharyngeal or transitional populations, C7 and C14 showed similar enrichment for OCs containing *Hmgb* genes (OC12, OC18, OC24, OC26) and immature mesenchymal markers (e.g. *Twist1* and *Prrx2* in OC100) (Figure5b, c, Table S9). Cushion-associated clusters C10 and C22 also shared similar OC profiles, though with reduced enrichment for *Hox*-containing OC45 and increased enrichment for *Tbx20*/*Gata4*-containing OC25, consistent with *Hox*-downregulation and corresponding changes in motif accessibility (Figure 4k, l, and 5b). The putative intermediate cluster C2 was characterized by OC25 and other *Tbx20*-containing OCs (OC86, OC91), bifurcating toward two major lineages: OC58^high^ valvular interstitial cells (C21; enriched for *Meox1, Sall3*, and *Scx*) and OC71^high^ subvalvular interstitial cells characterized by *Ebf2*, transitioning into coronary artery SMCs (C18 to C23) with distinct OC profiles (Figure S10b). Notably, coronary artery SMCs (C23) were characterized by enrichment of OCs containing *Stat3*, *Foxs1,* and/or *Epas1* (OC8, OC23, OC34) (Figure 5b and Figure S10b).

Clusters C7 and C14 are proposed progenitors of great artery SMCs (C0, C4, C27), aorticopulmonary septum (C16), and mediastinal mesenchyme including adipocytes (C6), all of which develop independently of the cardiac cushion. These populations exhibited relatively higher enrichment of *Hox*-containing OC45 compared to cushion-associated NCCs. Each component of great artery SMC lineage displayed region- and maturity-specific OC signatures (e.g., C27 vs. C4, and C0 vs. C27) (Figure S10b). In contrast to coronary SMCs, great artery SMCs showed higher enrichment for *Egr1*/*Mef2*-containing OCs (OC4, OC33) and lacked intermediate states marked by *Tbx20* and/or *Gata4*-containing OCs (Figure 5b and Figure S10b). The aorticopulmonary septum (C16) shared OC profiles with C2 but differed in its relatively higher enrichment of *Hox*-containing OCs (OC14, OC60) and lower enrichment of *Tbx20*-containing OCs (Figure 5b).

### Region-specific NCC gene expression is regulated by distinct Meis-binding motifs

To further investigate distal enhancers involved in TF-mediated, region-specific NCC lineage commitment, we developed a novel R package, CARTA (Connected Accessible Regions by input of the combination between Transcription factor and tArget), to construct *cis*-regulatory networks (CARTA-Net), extracting TF-binding motifs within enhancer-like regions that positively correlate with gene expression, are co-accessible with TSSs, and are highly conserved across mammals, using only TF-target combinations of interest as input (Figure 6a).

**Figure 6.**
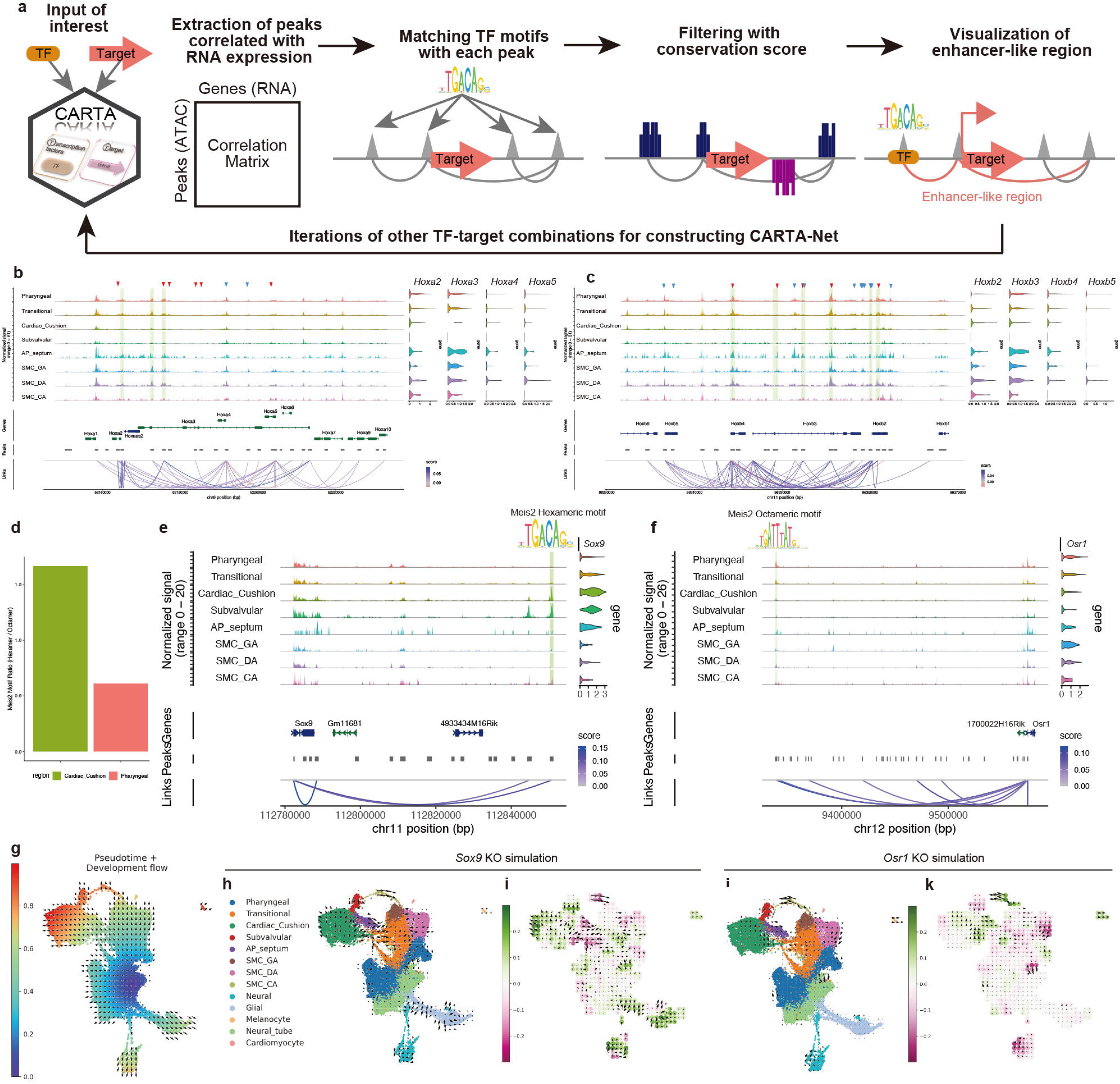
*Cis*-regulatory networks underlying cardiopharyngeal NCC development. (a) Workflow for CARTA-Net, which infers *cis*-regulatory networks from scMultiome datasets including scATAC-seq and scRNA-seq. (b, c) Coverage plots showing chromatin accessibility at *Hoxa* (b) and *Hoxb* (c) gene loci in NCC-derived mesenchymal subpopulations (left). Violin plots display corresponding *Hox* gene expression from scRNA-seq data (upper right). Locations of Meis-binding octameric motifs (MA1639.2 and MA1640.2) and hexameric motifs (MA0498.3 and MA0774.1) are indicated in red and blue arrowheads, respectively. Peaks differentially accessible in pharyngeal mesenchyme are highlighted in light green. (d) Comparison of the ratio of Meis-binding hexameric and octameric motifs in open chromatin peaks between pharyngeal and cardiac cushion mesenchyme. (e, f) Coverage plots of chromatin accessibility at *Sox9* (e) and *Osr1* (f) loci in NCC-derived mesenchymal subpopulations (left), with violin plots showing gene expression from scRNA-seq data (upper right). Differentially accessible peaks including Meis-binding motifs are highlighted in light green. (g) Pseudotime developmental flow of integrated NCC clusters from the neural tube, inferred using CellOracle and projected onto the UMAP space shown in Figure 4a. (h-k) *Sox9* (h, i) and *Osr1* (j, k) knockout simulations presented as altered differentiation vector flows (h, j) and perturbation scores, defined as the inner product between the simulated perturbation vectors and the original developmental flow (i, k). Green and magenta color bars indicate normal developmental flow and reverse flow induced by perturbation of the indicated genes, respectively.

To begin with, we examined chromatin accessibility at the *Hoxa* and *Hoxb* clusters and observed enrichment of MEIS-binding motifs within the accessible regions in pharyngeal and transitional mesenchymal NCCs, coinciding with high *Hox* gene expression (Figure 6b, c). Meis-binding octamer motifs were more enriched than hexamer motifs, especially in differentially accessible peaks of pharyngeal mesenchymal populations (Figure 6b, c). Recently, Kessler et al. identified a super-enhancer–rich region regulating anterior *Hoxa* collinearity via inter-topologically associating domain (TAD) interactions ^37^. This region contains two subdomains, HIRE1 and HIRE2, which are broadly accessible and marked by H3K27ac across cranial NCCs, including *Hox*-negative domains. However, inter-TAD interactions with Hoxa2 occur only in *Hoxa2*-expressing PA2 derivatives. Consistent with this, our data showed ATAC-seq peaks positively correlated with *Hoxa2* in pharyngeal and transitional mesenchyme (Figure S11a, b), but reduced in intracardiac mesenchyme except the AP septum. Most smooth muscle retained these peaks, whereas coronary artery smooth muscle showed reduced accessibility, in line with *Hox* expression. Conversely, intracardiac mesenchyme exhibited peaks negatively correlated with *Hoxa2* (Figure S11a, b). Motif analysis revealed enrichment of MEIS octamer motifs in pharyngeal/transitional mesenchyme, and Gata4-binding motifs in intracardiac mesenchyme (Figure S11c, d), consistent with gene regulatory changes during the pharyngeal-intracardiac NCC transition. Notably, these accessibility patterns differ from *Hox*-negative cranial NCCs, suggesting a distinct mechanism of *Hox* downregulation in intracardiac NCCs.

To investigate whether the functional transition of Meis TFs contributes to GRN diversification during cardiopharyngeal NCC differentiation, we further extended the CARTA-Net analysis (Table S10). Notably, hexameric and octameric Meis2-binding motifs were differentially associated with the regulation of target genes characteristic of cushion-associated and pharyngeal/cushion-independent mesenchymal subtypes, respectively (Figure 6d). These findings suggest that hexameric and octameric Meis2-binding motifs contribute to distinct region-specific transcriptional programs.

Among region-characterizing OCs, OC86, which characterized the cushion-associated intermediate cluster C2, contained *Meis2* and *Sox9* as well as *Tbx20* (Supplementary Table S9). Notably, a hexameric Meis2-binding motif (MA0774.1) was identified in the mesenchymal open chromatin region associated with the TSS of *Sox9* (Figure 6e and Table S10). In contrast, an octameric Meis2-binding motif (MA1640.1) was preferentially located in open chromatin regions associated with the TSS of genes like *Osr1*, characteristic of cushion-independent clusters (Figure 6f and Table S10). These contrasting expression patterns were validated through Xenium in situ analysis and further confirmed in the integrated UMAP (Figure 2n-u and Figure S3h, i).

Previous studies have implicated Meis2 and Sox9 in NCC development and the formation of cardiac outflow tract structures^38,39^. In contrast, Osr1 has been reported to suppress *Sox9* expression^40^, suggesting their antagonistic roles in NCC lineage diversification. To test this, we performed *in silico* gene perturbation using CellOracle^41^ on GRNs during lineage commitment. Depletion of *Sox9* in the GRN reversed the direction of differentiational flow from pharyngeal/transitional NCCs to intracardiac NCCs (Figure 6g-i). Conversely, *Osr1* depletion inhibited differentiation into great vessel SMCs while promoting differentiation into intracardiac NCCs (Figure 6g, j, k). Thus, Sox9 and Osr1 may function as antagonistic regulators in the lineage bifurcation from pharyngeal NCCs.

### A Meis-binding hexameric motif functions as a distal enhancer of *Sox9*

To further assess the functional role of Meis-binding motifs, we focused on the hexameric motif associated with *Sox9*. In ChIP-seq and ATAC-seq data of the embryonic heart provided by ENCODE, this region exhibited high accessibility and enrichment for H3K4 monomethylation with low levels of H3K27 acetylation (Figure 7a, Figure S12a), indicative of potential enhancer activity. In the developing heart, this region showed high accessibility in both *Tbx20*^high^ mesenchymal and epicardial cells, with high *Sox9* expression (Figure S12b). Luciferase assays using the O9-1 NCC line confirmed this, as an 888-bp fragment containing the hexameric Meis-binding motif significantly increased luciferase activity, while deletion of the hexameric Meis motif markedly attenuated this effect (Figure 7b, c). In vivo enhancer activity was further validated using *Sox9*-Enhancer-LacZ reporter mice, in which the same 888-bp fragment drove LacZ expression in NCC-derived cushion mesenchyme and also in the epicardium, both sites of robust *Sox9* expression (Figure 7d-h). Enhancer activity in the epicardium corresponds to *Sox9* expression and an open chromatin peak at the putative distal enhancer region in clusters 22 and 5 in Figure 1d, which represent *Wt1*^high^ epicardial cells and intracardiac mesenchyme likely including *Wt1*^low^ epicardial EMT derivatives, respectively (Figure S13). These results support the role of this hexameric Meis-binding motif-containing region as a distal enhancer of *Sox9* in cushion NCCs and other *Sox9* expressing cells.

**Figure 7.**
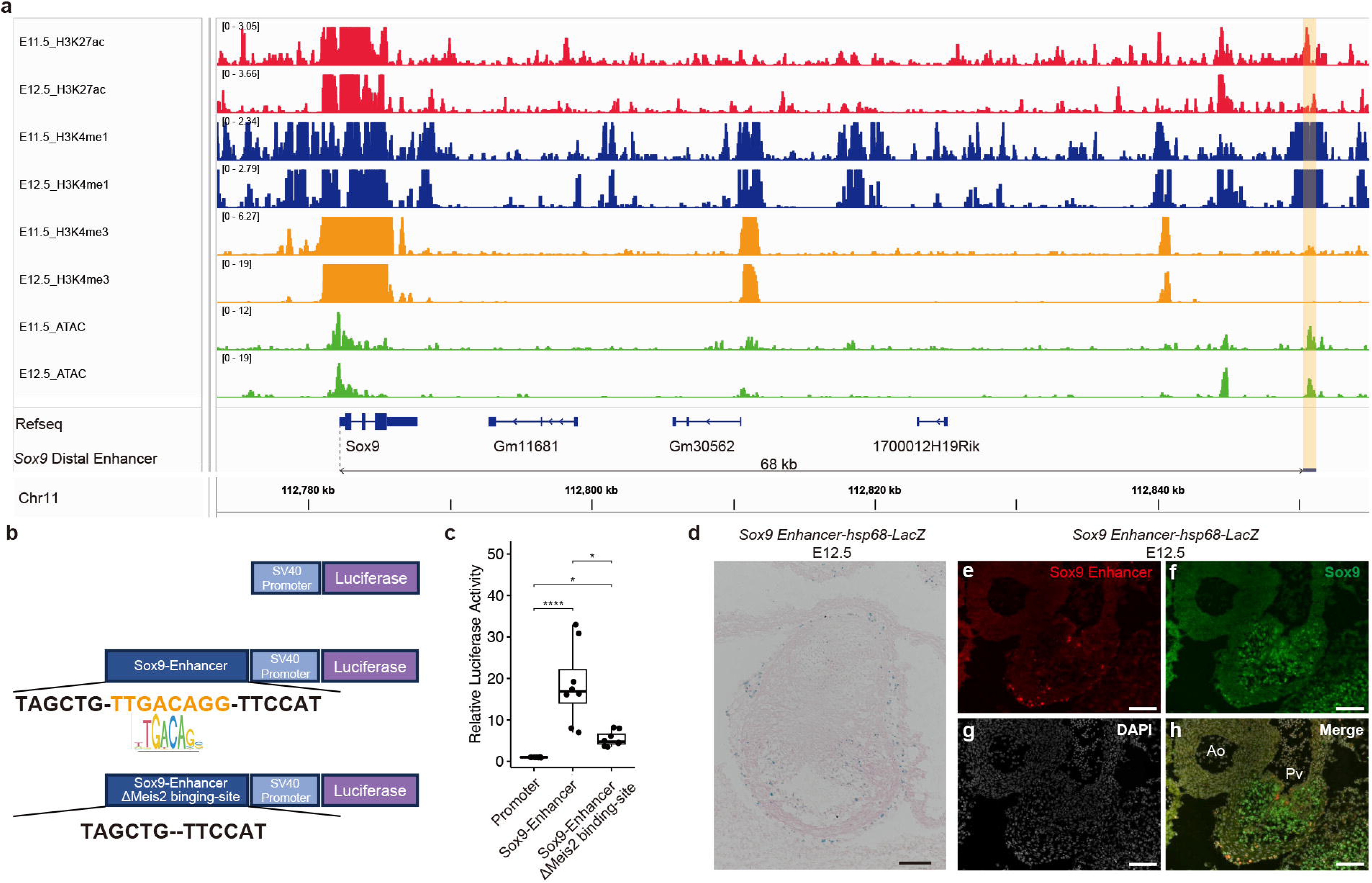
A potential *Sox9* enhancer region containing a hexameric Meis-binding motif. (a) ChIP-seq and ATAC-seq data of the embryonic heart from ENCODE, accessed via the UCSC Genome Browser (http://genome.ucsc.edu/). The candidate distal enhancer region is highlighted in orange. (b, c) Luciferase reporter assay using constructs containing the 888-bp fragment with either an intact or deleted hexameric Meis-binding motif (b), tested in the O9-1 NCC line (c). *P<0.05, ****P<0.0001. (d) X-gal staining forβ-galactosidase activity in the outflow tract cushion of *Sox9*-Enhancer-LacZ reporter mouse embryos at E12.5. Scale bar, 100 μm. (e-h) SPiDER β-Gal staining (e), Sox9 immunostaining (f), and DAPI nuclear staining (g) in the cushion region of E12.5 *Sox9*-Enhancer-LacZ embryos. Merged image shown in (h). Scale bars, 100 μm.

### Characterization of *Sox9*^high^/*Scx*^high^ NCC population and their lineages in the developing heart

OC86, a GRN module characterizing cushion-associated NCCs, includes *Scx*, a marker of tendon/ligament-related skeletogenic progenitors that acts alongside *Sox9* (Supplementary Table9), leading us to speculate that a similar *Sox9*^high^/*Scx*^high^ population may function as progenitors of cushion-associated NCC derivatives. In the integrated UMAP, *Sox9*^high^/*Scx*^high^ NCCs were enriched in C2, C10, C14 and C16.

To spatiotemporally identify this population, we used *Sox9EGFP* and *ScxTomato* mice, which allow visualization of *Sox9* and *Scx* expression, respectively (Figure 8a-c). At E12.5, EGFP signals marking *Sox9* expression were observed in the outflow tract cushion tissue, where αSMA was expressed at a lower level than the surrounding muscular tissues (Figure 8a). Within this *Sox9*-positive domain, *Scx*-tdTomato-expressing cells formed an aggregation, corresponding to NCC-derived mesenchymal condensation (Figure 8d). From E14.5 to E17.5, *Sox9* and *Scx* expression persisted in outflow cushion-derived semilunar valve tissues, especially in the fibrous interleaflet triangles, although expression levels varied (Figure 8b, c). By contrast, Myh11-labeled coronary artery SMCs, also derived from NCCs, lacked *Sox9* and *Scx* expression (Figure 8c).

**Figure 8.**
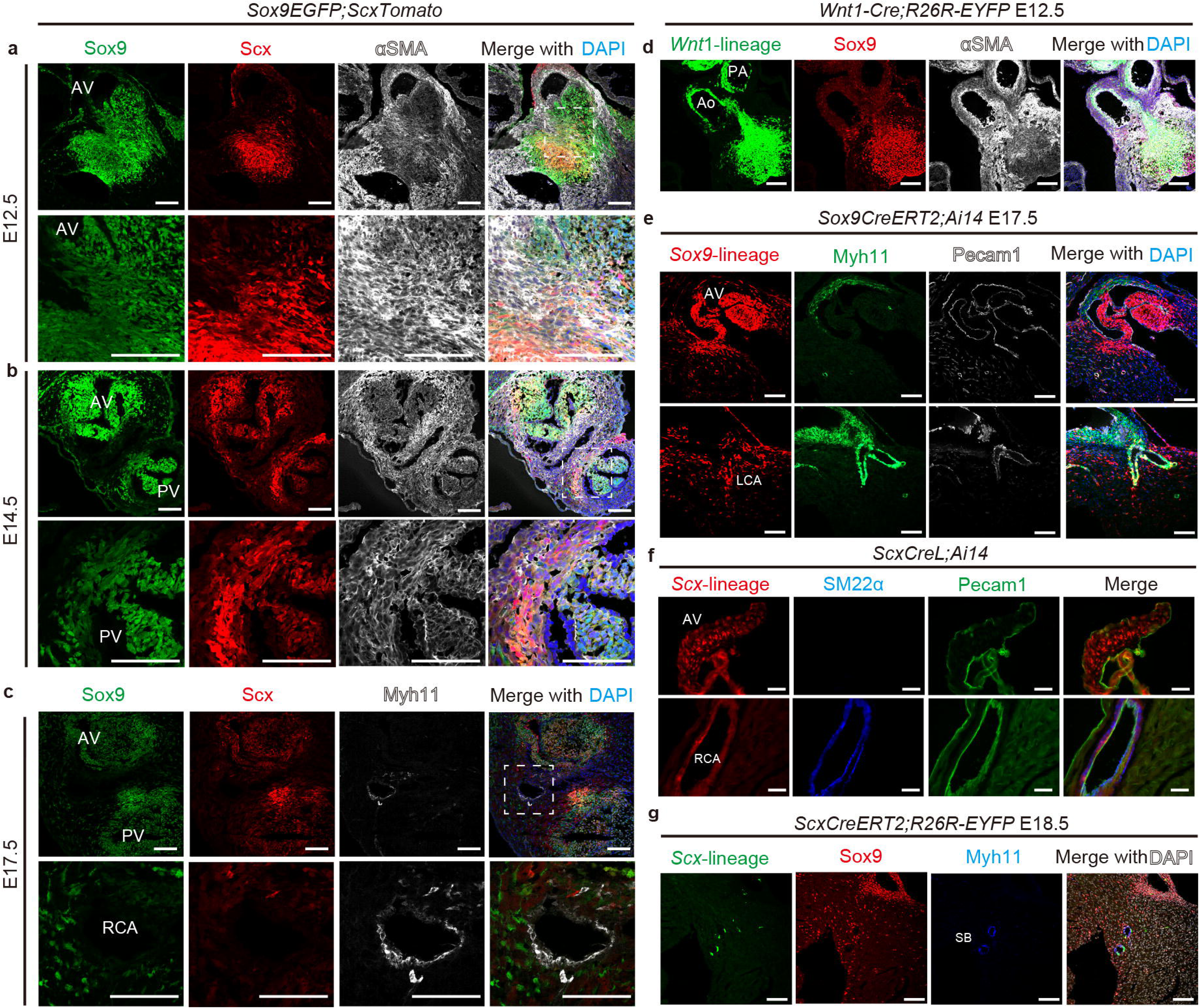
*Sox9*^high^/*Scx*^high^ cells and their descendants in the developing heart. (a-c) Immunostaining for Sox9-EGFP and Scx-tdTomato with co-staining for αSMA (a, b) or Myh11 (c) in *Sox9-EGFP;Scx-tdTomato* mice at E12.5 (a), E14.5 (b), and E17.5 (c). Boxed regions are magnified in lower panels. (d) Immunostaining for EYFP (*Wnt1*-lineage), Sox9, and αSMA in *Wnt1-Cre;R26-EYFP* mice at E12.5. (e) Immunostaining for tdTomato (*Sox9*-lineage), Myh11, and Pecam1 in E17.5 *Sox9-CreERT2;Ai14* mice treated with tamoxifen at E11.5. (f) Immunostaining for tdTomato (*Scx*-lineage), SM22α, and Pecam1 in *Scx-CreL;Ai14* mice at 3 months. (g) Immunostaining for EYFP (*Scx*-lineage), Sox9, and Myh11 in E18.5 *Scx-CreERT2;R26R-EYFP* mice treated with tamoxifen at E12.5. Ao, aorta; AV, aortic valve; LCA, left coronary artery; PA, pulmonary artery; PV, pulmonary valve; RCA, right coronary artery; SB septal branch of coronary artery. Nuclei are counterstained with DAPI. Scale bars, 50 μm.

To determine whether valve leaflets and coronary artery SMCs differentiate through a *Sox9*^high^/*Scx*^high^ intermediate state, we employed Cre reporter mice. In *Sox9CreERT2;Ai14* mice treated with tamoxifen at E11.5, Cre-mediated recombination was broadly detected in the semilunar valve leaflets, coronary artery SMCs, and surrounding mesenchymal cells at E17.5 (Figure 8e). Similarly, *ScxCreL;Ai14* mice exhibited labeling in both semilunar valve leaflets and coronary artery SMCs (Figure 8f). Moreover, tamoxifen administration at E12.5 in *ScxCreERT2;R26R-EYFP* mice resulted in EYFP expression in coronary artery SMCs (Figure 8g).

These findings collectively support the existence of *Sox9*^high^/*Scx*^high^ intermediate population in and around the mesenchymal condensation, from which semilunar valve tissues and coronary artery SMCs of NCC origin are derived. Together with our previous report that proximal coronary artery SMCs originate from preotic rather than postotic NCCs^8^, these results suggest that the intermediate population contributing to coronary artery SMCs likely represents a subset of *Hox*-downregulated intracardiac NCCs corresponding to clusters C2 and C10.

## Discussion

In this study, we present a comprehensive map of cardiopharyngeal NCC populations by integrating single-cell and spatial transcriptomics with publicly available datasets. This map bridges early pharyngeal NCCs and late-stage cardiac derivatives, both through and independent of the outflow tract cushion. Cushion-associated intracardiac NCCs showed distinct *Hox* gene downregulation, coinciding with a shift in Meis transcription factor binding from Hox-dependent (octameric motifs) to non-Hox (hexameric motifs) partners. GRN analysis linked these motif types to region-specific targets such as *Sox9* and *Osr1*, defining distinct regulatory modules. We further identified distinct intracardiac mesenchymal populations, including aorticopulmonary septum, subvalvular mesenchyme, and coronary artery SMC–related intermediates, thereby refining the developmental trajectories of cardiac NCC derivatives. These findings are summarized in Figure 9.

**Figure 9.**
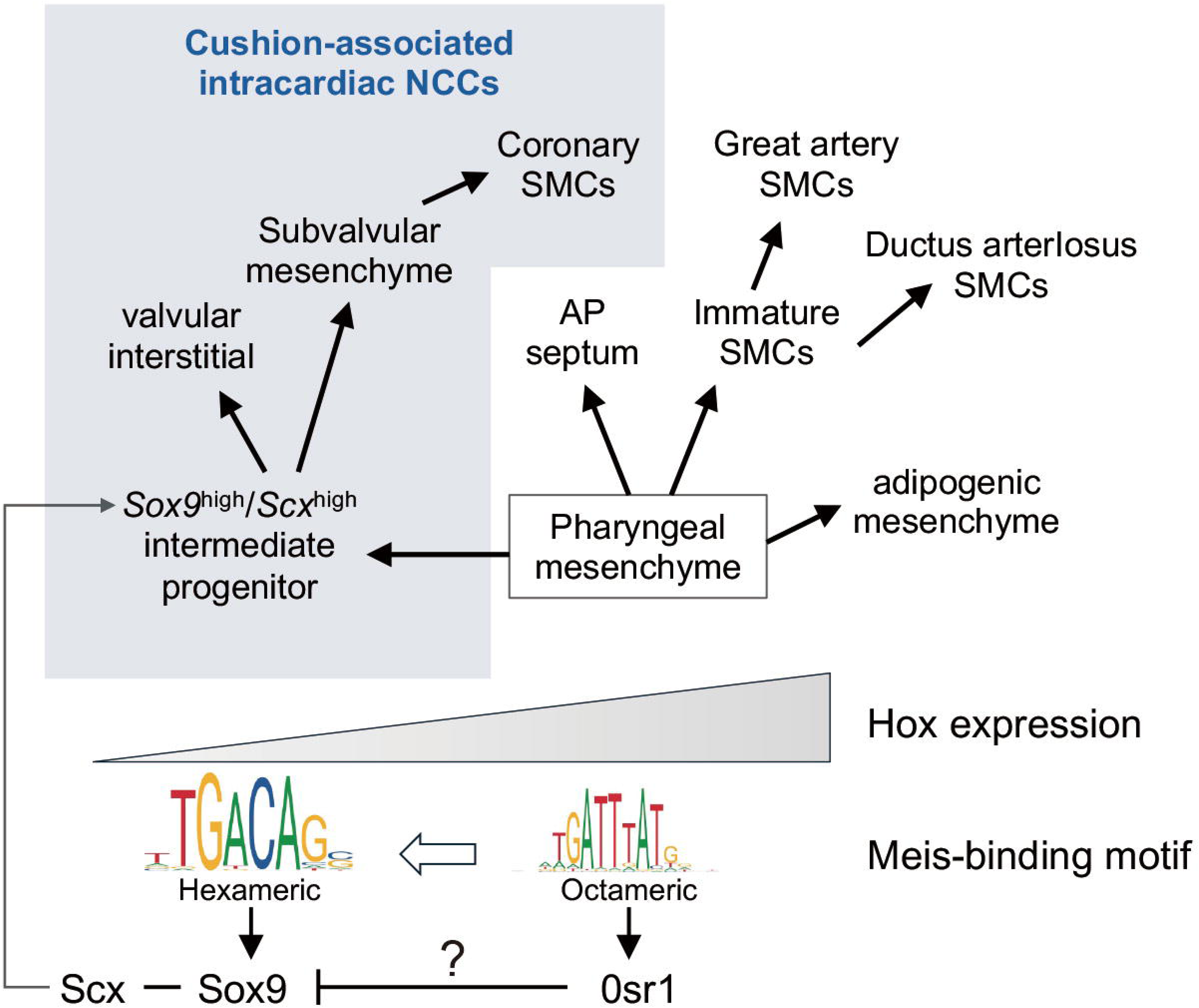
Summary schematic illustrating the diversification of cardiopharyngeal NCC lineages orchestrated by a regional genetic switch involving a Hox–Meis relay.

Our integrated map incorporates previously published lineage analyses of cardiac NCCs at early and late stages^13,14^, providing continuity through complementary single-cell and spatial transcriptomic data. The present study further extends these datasets by resolving the heterogeneity of intracardiac mesenchymal populations and their lineage relationships. For example, the cell population identified by De Bono et al. as outflow smooth muscle^13^ corresponds in our dataset to early intracardiac mesenchymal clusters expressing immature SMC markers, which subsequently diverged into multiple derivatives including coronary artery SMCs. In addition, we identified distinct SMC populations corresponding to great artery SMCs, ductus arteriosus SMCs, and coronary artery SMCs, each characterized by unique molecular signatures such as *Sost*, *Tfap2b*/*Ptger4*, and *Reln*/*Gja4*, respectively. The continuity between intracardiac mesenchyme and coronary artery SMCs through a pericyte-like intermediate state is consistent with previous developmental studies showing that proximal coronary artery SMCs originate from preotic NCCs and may arise through pericyte intermediates^8,23^. Together, these findings provide a refined framework for understanding the diversification of cardiac NCC derivatives during outflow tract remodeling.

The regional identities of pharyngeal NCCs that contribute to cardiac development are established by *Hox* genes and their associated regulatory networks. Genetic studies have demonstrated essential roles for the anterior *Hox* genes in patterning the pharyngeal arch artery system and semilunar valve structures. Loss of *Hoxa1* and *Hoxb1* results in severe defects in pharyngeal arch artery development^42^, whereas ectopic or sustained expression of *Hoxb1* in NCCs disrupts cardiovascular morphogenesis and causes malformations of the great arteries and semilunar valves^43^. Likewise, *Hoxa3* contributes to proper patterning of the pharyngeal arch region and its NCC-derived derivatives^44,45^. These findings underscore the importance of precise spatial and temporal regulation of *Hox* genes during cardiovascular development. Among pharyngeal NCCs contributing to cardiac development, cushion-independent NCC derivatives (great artery SMCs and the aorticopulmonary septum) retain their origin-specific *Hox*-codes. In contrast, cushion-associated NCC derivatives (coronary artery SMCs and valvular/subvalvular interstitial cells) downregulate *Hox* expression and transition toward region-specific GRNs involving TFs such as *Tbx20* and *Gata4*, whose expression is known to be induced by BMP signaling in cardiomyocytes^46,47^. *Bmp2* and *Bmp4* are expressed in the regions of the pericardial reflection traversed by NCCs en route to the cardiac cushion^48^. Together, these observations suggest that appropriate repression of *Hox* programs, coupled with activation of cardiac-specific regulatory networks, is required for normal differentiation of cushion-associated NCC derivatives.

The transcriptional transition accompanying entry into the cardiac cushion was associated with extensive chromatin remodeling. Accessibility of *Hoxa*-associated regulatory regions and enrichment of Meis-binding octamer motifs were maintained in pharyngeal and transitional mesenchyme but markedly reduced in intracardiac mesenchyme, except for the aorticopulmonary septum. Conversely, intracardiac mesenchyme showed enrichment of Gata4-binding motifs and *Hoxa2*-negatively correlated accessible regions, consistent with a shift toward cardiac-specific regulatory programs. These findings align with the recent report by Kessler et al.^37^ describing inter-TAD regulation of anterior *Hoxa* genes in cranial NCCs, but suggest that intracardiac NCCs undergo a distinct mode of *Hox* silencing associated with lineage-specific chromatin reorganization. Together with the Xenium observation that *Tbx20* expression in cardiac tissues was distinctly demarcated from the pharyngeal regions, these findings suggest that local cardiac signaling environments, potentially including BMP signaling, may play a critical role in the gene regulatory switch in cushion-associated NCC differentiation.

Concomitant with *Hox* gene downregulation upon entering the cardiac cushion, NCCs undergo a shift in Meis protein DNA-binding preferences, which typically function as cofactors of Hox and Pbx in early mesenchyme, from octameric to hexameric motifs, despite stable Meis1/2 expression levels. This switch was accompanied by increased expression and motif accessibility of cardiac-specific TFs such as Gata4 and Tbx20. Recently, Darieva et al. demonstrate that Meis proteins interact with Gata4 at cardiac-specific enhancers in a cardiomyocyte differentiation context^49^. Taken together, these findings suggest that Meis proteins adapt their transcriptional partners in a region-specific manner, enabling context-dependent gene expression. This combinatorial TF switching may represent a regulatory logic for establishing cushion-associated NCC identity and directing their differentiation trajectories.

To further investigate these mechanisms, we developed the CARTA package to identify candidate TF-binding enhancers driving GRNs in distinct NCC subpopulations. Using this approach, we identified a putative distal enhancer of *Sox9*, containing a hexameric Meis2-binding motif. This region was enriched for H3K4 monomethylation in the embryonic mouse heart, aligning with Darieva et al.’s report that MEIS proteins recruit the H3K4 monomethyltransferase KMT2D, to initiate lineage-specific enhancer commissioning^49^. The enhancer activity of this region was validated through both *in vitro* and *in vivo* assays. These results support a model in which MEIS TFs, through motif-specific enhancer binding and cofactor recruitment, orchestrate GRN divergence and fate specification in cardiac NCCs.

We also identified a *Sox9*^high^/*Scx*^high^ intermediate population involved in intracardiac NCC differentiation into coronary artery SMCs and semilunar valve interstitial cells. This population shares transcriptional and morphological characteristics with skeletogenic progenitors, including the formation of mesenchymal condensations^50,51^. This suggests the presence of shared GRNs underlying both differentiation programs. Previous reports have shown that knockout of *Ets1*, *Fn1*, or *Adam19* in mice results in NCC-derived ectopic cartilage formation in the heart^52–54^, implicating these genes in the repression of skeletogenic programs in intracardiac NCCs. At later developmental stages, *Sox9* and *Scx* expression persists at the bases of the semilunar valve leaflets. In skeletal tissue development, these genes are co-expressed in teno-chondrogenic progenitors that form the tendon-bone attachment unit^55,56^. By analogy, *Sox9*^high^/*Scx*^high^ NCCs at the base of semilunar valves may form a structural attachment unit linking cushion tissues to valvular leaflets^57^.

Overall, this study proposes a new framework for understanding cardiac NCCs heterogeneity based on developmental route, *Hox*-code retention, and region-specific regulatory programs. Importantly, the developmental relationships and differentiation pathways described here are inferred from integrated computational analyses, including transcriptomic similarity, UMAP connectivity, and RNA velocity, rather than direct lineage-tracing experiments. Within this framework, our findings suggest distinct differentiation trajectories leading to great artery, ductus arteriosus, and coronary artery SMCs, as well as the aorticopulmonary septum and valvular/subvalvular mesenchyme. Furthermore, we identify an intermediate progenitor population within the cushion-associated NCC lineage that differentiates into coronary artery SMCs and semilunar valve components. Notably, both the coronary artery and the aortic valve are highly susceptible to calcification, yet the developmental basis for this propensity remain poorly understood. The observed similarities and differences in GRNs between cardiac NCCs and skeletogenic progenitors may offer new insights into understanding the pathogenesis of calcification.

## Methods

### Animals

*Wnt1-Cre*^18^, *Rosa26-loxp-stop-loxp-EYFP* (*R26R-EYFP*)^58^, *Sox9EGFP*^59^, *Sox9CreERT2*^60^, *ScxCreL*^55^, *ScxCreERT2*^61^, *R26R-CAG-loxP-stop-loxP-tdTomato* (*Ai14*)^62^, and *ScxTomato*^61^ mice have been described previously. *Sox9* Enhancer*-hsp68-LacZ* transgenic mice with or without Meis2-binding site were generated by injecting the linearized *Sox9* distal enhancer*-hsp68* minimal promoter*-*nls*-LacZ* transgene into pronuclei of BDF1 fertilized eggs as described previously^63^. Mutant mice were maintained on a mixed C57BL/6J × ICR background. All animal experiments were approved by the Ethics Committee for Animal Experiments of The University of Tokyo, the Committee of Animal Experimentation of Hiroshima University, and Committee of Animal Experimentation of Ritsumeikan University. Mice were housed at 23±2□ with a relative humidity of 50-60% and light cycles with 12 hours light and 12 hours dark.

### Single-cell multiome (scRNA-seq and scATAC-seq) of cardiopharyngeal NCCs

Cardiopharyngeal tissues (E11.5 and E12.5) and cardiac outflow tract tissues (E14.5 and E17.5) were isolated from *Wnt1-Cre;R26R-EYFP* embryos. Tissues were dissociated by using 0.25 w/v% trypsin / 1 mmol/L EDTA-4Na solution (Wako) at 37°C, 15 minutes and neutralized with equal volume of DMEM (Wako) with 10% fetal bovine serum (FBS) (SIGMA). Cell suspensions were filtered through a 35 μm nylon mesh cell strainer (FALCON 352235) and kept on ice until cell sorting. Single-cell suspensions were stained with 7-AAD (BD Pharmingen) for 3 min at 4□, and EYFP-positive and -negative single cells were sorted using a FACSAria II or FACSMelody (BD Biosciences). The sorting strategy was as follows:

Step 1. All events were gated by forward scatter (FSC) and side scatter (SSC) including area (A), height (H), and width (W) to obtain FSC singlets and remove doublets.

Step 2. FSC singlets were gated for the 7-AAD negative fraction to isolate viable cells. Step 3. Viable cells were gated to isolate EYFP-positive NCCs or EYFP-negative non-NCCs.

Sorted cells were freshly processed or cryopreserved for the following procedure.

Frozen cells were thawed in DMEM with 10% FBS and nuclei were isolated using the protocol of Nuclei Isolation for Single Cell Multiome ATAC + Gene Expression Sequencing for low cell input (10x Genomics). Transposase-mediated insertion of adapters into open chromatin regions was followed by encapsulation into droplets with gel beads (GEMs) for scRNA-seq reverse transcription. cDNA and ATAC libraries were prepared using the Chromium Next GEM Single Cell Multiome ATAC + Gene Expression kit (10x Genomics) and sequenced on an Illumina NovaSeq 6000.

### Single-cell RNA sequencing (scRNA-seq) of cardiac NCCs with Fluidigm C1 system

EYFP-positive cardiac NCCs from *Wnt1-Cre;R26R-EYFP* mice at E11.5, E12.5, E14.5, and E17.5 were dissociated as above and captured using the Fluidigm C1 Medium-cell (10-17 μm cell diameter) integrated fluidic circuit (IFC) chips at 300 cells/μL. Each capture chamber was photographed (KEYENCE BZ-X710) to confirm single EYFP-positive cells. Single-cell cDNAs were prepared using SMART-Seq v4 Ultra Input Low RNA kit for the Fluidigm C1 System (Clontech), and quality was confirmed with the Agilent 2200 TapeStation and quantified by Qubit. (Thermo Fisher). High quality cDNAs were further subjected to the construction of sequencing libraries by using Nextera XT DNA Sample Preparation Kit (Illumina) and sequenced with 50 bp pair-end reads on the Illumina HiSeq 2500.

### Single-cell multiome data analysis

The sequence output FASTQ files were aligned to mouse (*Mus musculus*) genome reference (refdata-cellranger-arc-mm10-2020-A-2.0.0) provided by 10x Genomics adding EYFP sequence to produce gene expression counts by scRNA-seq and fragment counts by scATAC-seq in Cell Ranger ARC v2.0.0 pipeline (10x Genomics). Processing was performed using the SHIROKANE supercomputing resource provided by Human Genome Center (the Univ. of Tokyo). Downstream data analysis was performed in R version 4.4.0 and Python 3.8. The gene expression and fragment counts were converted into Seurat objects by using Seurat version 5.1.0 and Signac version 1.4.0 R package^64,65^.

The percentage of mitochondrial genes was calculated by the PercentageFeatureSet function. For the further QC filtering, SoupX ^66^ was used for the removal of cell free RNA contamination in droplet-based scRNA-seq data. Multiplets were removed using DoubletFinder ^67^.

Filtering thresholds were: RNA counts >3000 and <50,000; ATAC counts >3000 and <500,000; TSS enrichment >2; nucleosome signal <4; mitochondrial content <25%. The cell cycle score was calculated by using the CellCycleScoring function. Then, the scRNA-seq dataset was subjected to dimensional reduction into a two-dimensional UMAP space with RPCA integration to reduce technical batch effects. Cell clustering was performed by constructing a KNN graph based on the Euclidean distance provided by principal component analysis and the Louvain algorithm. Differential expressed genes (DEGs) and Differential accessible peaks (DAPs) were calculated by Wilcoxon rank-sum test. TF binding motifs in highly accessible chromatin regions were analyzed by ChromVAR^68^. These expression profiles were visualized by FeaturePlot function from Seurat. RNA Velocity analysis was performed by velocyto.py version 0.17and scVelo version 0.3.2.^69,70^ Pseudotime was set as centroid of the immature cell clusters such as C1, C3, C19 (neural tube). The NCC lineage diversification flow focused on the transitional state was analyzed by CellRank2 version 2.0.7^30^. The association test was used to detect differential expressed genes along the pseudotime by Slingshot version 2.12.0^29^. CellOracle^41^ was used for gene perturbation analysis.

### Fluidigm C1 scRNA-seq data analysis

The sequence output FASTQ files were aligned to indexed mouse (*Mus musculus*) genome reference (GRCm38/mm10) with EYFP sequences by using HISAT2 software version 2.1.0. Gene expression counts were calculated with the featureCounts function from the Rsubread version 1.34.7 package using R and converted into Seurat objects. The percentage of mitochondrial genes against detected genes per cell was calculated using the top 200 mitochondrial genes from the MitoCarta 2.0 database. Cells with feature RNA >2000 and mitochondrial content <10% were used for further analysis. The cell cycle score was calculated as above. The projection of Fluidigm C1 scRNA-seq data onto the UMAP of integrated scRNA-seq datasets was performed by anchor-based transfer using FindTransferAnchors and MapQuery functions in Seurat.

### Estimation of GRNs

We used SCENIC^35^ version 1.1.2.2 R package to extract the gene combinations inferred by gene regulatory motifs and SiGN-BN NNSR version 0.16.6^34^ to construct GRNs on the SHIROKANE supercomputer. The Fluidigm C1 scRNA-seq expression count data were converted into transcripts per million (TPM) to minimize mapping biases associated with full-length scRNA-seq. TPM values were analyzed using the SCENIC pipeline. Gene filtering was performed default geneFiltering function in SCENIC. Spearman correlation coefficients were then calculated using runCorrelation function and gene pairs with correlation values >0.03 (default) were used to construct TF–gene co-expression modules by using runGenie3 function.

Regulons, defined as TF–gene pairs supported by TF-binding motifs, were subsequently inferred using RcisTarget with motif databases: mm9-500bp-upstream-7species.mc9nr.feather and mm9-tss-centered-10kb-7species.mc9nr.feather. Subsequently, we applied the SiGN-BN NNSR algorithm to TPM expression data, filtered to include only genes identified by the Seurat FindAllMarkers function with the parameter of only.pos = FALSE, to construct a GRN based on nonparametric Bayesian estimation. SiGN-BN NNSR was run at the following parameters: m (maximum number of parents that per gene) = 1000; T(number of iterations for subnetwork estimation using the neighbor node sampling and repeat algorithm) = 1000000; skel-type = TXT--skel; parent–child edge combinations were restricted to those supported by regulons identified in the SCENIC analysis, further filtered by Genie3Weight > 0.004.

### Community analysis of GRNs

TF-TF subnetworks were extracted from GRNs and analyzed using the linkcomm R package^36^ to detect overlapping communities. The set of first-edge connections from TFs within each community were defined as an “overall community” (OC). The enrichment score of each OC per cell was calculated in three steps: Step1; Normalized expression counts for each gene in a community were converted into z-scores. Step2; For each cell, the z-scores of genes belonging to a given community were averaged. Step3; These cell-level scores were then averaged across cells within each cluster to obtain a cluster-level OC score. Spatial community detection was performed in a similar manner. OC enrichment scores were visualized as heatmaps using the pheatmap R package. To extract key OCs in each cell cluster, Random Forest classifier provided in caret R package was used. The expression matrix was divided into training and testing sets in a 7:3 ratio.

### Development of CARTA

CARTA was developed in R to construct *cis*-regulatory TF network (CARTA-Net) based on user-defined TF–target gene combinations using single-cell multiome data (scRNA-seq and scATAC-seq). The workflow consists of the following steps. Step1 (Feature Filtering); TFs, target genes, and peaks are filtered based on average log2 fold change, *p*-value, and average expression across cell clusters. This filtering step reduces computational burden by limiting the analysis to cluster-specific DEGs, TFs, and associated peaks. Step2 (Correlation Analysis); Pearson correlation coefficients were calculated between (i) TF and target gene expression, and (ii) target gene expression and peak accessibility (peak counts). Peaks located within ±500 kb from the transcription start site (TSS) of the target gene are considered by default. Only positive correlations are used in subsequent steps to identify putative enhancer regions. Step3 (Motif Matching); TF binding motifs are identified in correlated peaks by motifmatchr version 1.14.0 R package^71^, based on vertebrate motif catalogs in JASPAR2020^72^. The output includes the genomic loci and sequence of matched motifs. Step4 (Conservation Filtering); these motifs are filtered based on sequence conservation across Euarchontoglires with phastCon score^73^. We used the conservation score bigwig file based on the mm10, mm10.60way.phastCons60wayEuarchontoGlire.bw, to import for R with rtracklayer package. The average conserved score with more than 0.8 was used for the filtering. Step5 (Visualization); Visualization of enhancer-like regions including TF-binding motifs as the coverage plot. Enhancer-like regions containing TF-binding motifs are visualized as coverage plots. The entire workflow was iteratively applied for each specified TF–target gene pair to construct the final CARTA-Net.

### Xenium spatial transcriptome of embryonic sections

Unfixed mouse embryos at E11.5 and E12.5 were directly embedded in OCT compound (Sakura Finetek) and stored at –80□ until sectioning. These samples were cut at a thickness of 10 μm using cryostat (TheremoFisher) and mounted onto Xenium slides (10x Genomics). Multiplexed *in situ* hybridization was then performed using the Xenium Analyzer (10x Genomics) according to the manufacturer’s protocol. A customized panel of 334 target probes was used for this experiment. After data acquisition with the Xenium Analyzer, the slides were stained with hematoxylin and eosin (HE), and images were captured using the KEYENCE BZ-X710 microscope.

### Xenium spatial transcriptome data analysis

Spatial transcriptomic data output from the Xenium Analyzer were aligned with the corresponding HE-stained images, which were converted to OME.tif using QuPath version 0.5.0. These combined spatial transcriptome data was visualized by Xenium Explorer version 3.2.0. These data were processed with Seurat v5.1.0 to integrate each section to generate a unified UMAP by Sketch integration. DEGs were calculated by the same way described above. Tangram^17^ was used to impute the expression of genes not included in the custom probe set, based on the scRNA-seq dataset. Decomposition of cell clusters of scRNA-seq was performed by RCTD^19^ to map them onto the Xenium dataset. The pooled scRNA-seq data of NCCs or non-NCCs at E11.5 and E12.5 were used for RCTD. Putative NCCs were identified through the following procedure:

Step 1. EYFP expression was estimated across 39 cell types in the Xenium dataset by integration with the single-cell multiome dataset (including both NCCs and non-NCCs) using Tangram.

Step 2. Xenium clusters enriched for EYFP expression, defined as clusters whose mean estimated EYFP level exceeded the threshold corresponding to the 65th percentile across all spots, and consistent with known neural crest derivatives were extracted as putative NCC populations.

Step 3. For each spot within these EYFP-enriched Xenium clusters, RCTD was used to estimate the corresponding multiome cluster identity.

### Visium spatial transcriptome of embryonic hearts

Embryonic hearts from *Wnt1-Cre;R26R-EYFP* mice at E14.5 and 17.5 were embedded in a chilled OCT compound and immediately frozen on the metal plate in a liquid nitrogen bath. Frozen samples were cut at the thickness of 10 μm in the cryostat and placed on the Visium Spatial Tissue Optimization Slide to confirm the time to lyse sections and on the Visium Spatial Gene Expression Slide to get RNA-seq data. The lysis time to extract RNA from the section was determined by the results of Visium Tissue Optimization Kits (10x Genomics). Then, the cryosections on the Visium Spatial Gene Expression Slide were stained with HE and captured images with KEYENCE BZ-X710. After imaging, the sections were lysed in 24 and 18 minutes at E14.5 and 17.5, respectively, with Visium Spatial Gene Expression Reagent Kits (10x Genomics). mRNAs captured on the slides were reverse-transcribed into cDNA and used to construct sequencing libraries following the manufacturer’s protocol. Each of the libraries was sequenced on Illumina NovaSeq6000.

### Visium spatial transcriptome data analysis

FASTQ files from four datasets (E14.5_1, E14.5_2, E17.5_1, E17.5_2) were aligned to indexed mouse (*Mus musculus*) genome reference (GRCm38/mm10) with an additional EYFP sequences and gene expression counts were calculated by using Space Ranger version 1.1.0 (10x Genomics). The gene expression counts were converted into Seurat objects and region of interest were extracted to exclude the noise. After normalization with the SCTransform function, datasets were merged by developmental stage into E14.5 and E17.5 groups. Dimensionality reduction was performed using UMAP, and spot clustering was carried out as described above. Differentially expressed genes (DEGs) across spatial clusters were identified using the Wilcoxon rank-sum test, with a log fold-change threshold of >0.25. To identify cardiac NCCs in the tissue sections, spatial gene expression spots containing at least one mapped EYFP transcript were selected for further analysis. Spatial mapping of scRNA-seq data onto Visium sections was performed using only E14.5 and E17.5 datasets and the RCTD algorithm, as described above.

### Analysis of public datasets

We analyzed the publicly available scRNA-seq datasets of mouse NCCs deposited in Sequence Read Archive with the accession number: PRJNA562135^14^ and Gene Expression Omnibus with accession number: GSE210521^13^. The first dataset was aligned to mm10 reference genome using Cell Ranger ARC v2.0.0 pipeline (10x Genomics), and the second was processed using10x Genomics Cloud Analysis. Gene expression matrix data were converted into Seurat objects and analyzed as described above.

### Lineage tracing experiment

For lineage tracing of Sox9-positive cells in *Sox9-CreERT2;R26R-tdTomato* mice at E11.5, Pregnant female mice were injected with 2.5 mg of 4-hydroxytamoxifen (Sigma-Aldrich). For lineage tracing of Scx-positive cells in *ScxCreERT2;R26R-EYFP* mice at E12.5, tamoxifen (Sigma-Aldrich) was administered orally in corn oil (Sigma-Aldrich) at a dose of 0.1mg/g body weight. Embryos were sampled at appropriate stages.

### Immunohistochemistry

Embryos were fixed in 4% paraformaldehyde phosphate buffer solution (Nacalai Tesque) for 3 hours at 4□. Fixed embryos were embedded in OCT compound (Sakura Finetek) through stepwise sucrose substitution and stored at –20□ until sectioning. Samples were cut into 10 μm sections using cryostat (TheremoFisher). For immunohistochemistry, frozen sections were rinsed with phosphate-buffered saline (PBS) and permeabilized with 0.3 % Triton-X100 in PBS. Blocking was performed using 3% bovine serum albumin (BSA) in PBS. Sections were incubated with primary antibodies diluted in 3% BSA overnight at 4□. After washing with PBS, secondary antibodies diluted in 3% BSA were applied for 2 hours at room temperature. Antibodies were: SM22α (ab14106, Abcam, 1:800), αSMA (A2547, Sigma, 1:1000), GFP (04404-26, Nacalai Tesque, 1:1000), mCherrry (AB0040-200, SICGEN, 1:500), smooth muscle myosin heavy chain 11 (ab53219, Abcam, 1:200), Dlk1 (AF1144, R&D systems, 1:500), Sox9 (AF3075, R&D Systems, 1:500), Pecam1 (553370, BD Pharmingen, 1:200), and Alexa Fluor (488, 555 and 647)-conjugated secondary antibodies (Abcam, 1:200). Nuclei were stained with 4’,6-diamidino-2-phenylindole dihydrochloride (DAPI) in PBS (1:1000). Immunofluorescence images were captured by a Nikon C2 confocal microscope and BZ-X710 and BZ-X810 microscopes.

### LacZ staining

For whole-mount LacZ staining, embryos were fixed in 4% paraformaldehyde phosphate buffer solution 15 min at 4□ and rinsed twice in PBS containing 2 mM MgCl_2_, 10 mM EGTA, 0.02% NP-40, and 0.01% sodium deoxycholate for 10 min at 4□. The buffer was replaced with the detergent rinse buffer (80mM K_2_HPO_4_, 5 mM KH_2_PO_4_, 2 mM MgCl_2_, 0.02% NP40, and 0.01% sodium deoxycholate) for 10 min at 4□. Embryos were stained with the same buffer containing 10 mM K_3_(Fe(CN)_6_), 10 mM K_4_(Fe(CN)_6_), and 1 mg/ mL X-gal overnight at 37□. For section LacZ staining, fresh embryos were embedded in OCT compound (Sakura Finetek) and cut into 10 μm sections using cryostat. Sections were stocked at -80□ until staining. Sections were air dried at room temperature for 10 min and fixed with 4% paraformaldehyde phosphate buffer solution for 10 min at 4□. After rinsing with PBS twice, LacZ staining was performed as described above.

For fluorescence imaging, air dried sections were fixed as described above and rinsed with PBS three times for 3 min at 4□. SPiDER-βgal (Dojindo, SG02) was diluted in PBS to 1 μM, applied to sections, and incubated for 15 min at 37□. After incubation, sections were washed with PBS and co-immunostained as described above. Fluorescence images were acquired using a KEYENCE BZ-X710 microscope.

### Cell culture

Cell culture of the O9-1 neural crest cell line (Merck, SCC049) was conducted as described previously^74^. Culture dishes were coated with 20 μg/mL fibronectin. To maintain cells in an undifferentiated state, conditioned medium derived from STO cells (RIKEN BioResource Research Center) was supplemented with 10^3^ units/mL mouse LIF (Nacalai Tesque, NU0012-1) and 25□ng/mL mouse bFGF (BioLegend, 579604).

### Luciferase assay

Transfection was performed using Lipofectamine3000 Reagent (Invitrogen) with pGL3-luciferase reporter plasmids, including pGL3-Promoter, pGL3-Control, or pGL3-Enhancer (Promega) containing the *Sox9* distal enhancer-like region (chr11-112850248-112851135). The pGL3 vectors were co-transfected with Renilla luciferase vector phRL-TK as an internal control. Two days after transfection, cell lysates were collected using the Dual-Luciferase Reporter Assay System (E1960, Promega), and luciferase activities were measured using a Lumat LB 9507 luminometer (Berthold Technologies). Firefly luciferase activity was normalized to Renilla luciferase activity.

## Data availability

The 10x Genomics scMultiome data have been deposited in DDBJ Sequence Read Archive (DRA) under accession code: DRA015815, DRA015814, DRA014005, and DRA012897. Visium spatial transcriptome data was under DRA010734 and DDBJ Genomic Expression Archive (GEA) under accession code: GEAD-430. Xenium spatial transcriptome data have been deposited under accession code: GEAD-686 and A-GEAD-2. The Fluidigm C1 scRNA-seq data have been deposited in the Gene Expression Omnibus under accession code GSE201417.

## Code availability

An open-source R package of CARTA is available at GitHub (https://github.com/iaki-dev/CARTA). The scripts used for the analyses of 10x Genomics scMultiome, Visium, Xenium, Fluidigm C1 scRNA-seq, and public data are available at https://github.com/iaki-dev/Iwase_etal_2025.

## Acknowledgments

We thank Yoshinori Tamada (Hirosaki University) for technical guidance and the use of SiGN-BN NNSR on the super-computing resource provided by Human Genome Center (the University of Tokyo), Kiyomi Imamura, Kazumi Abe, Etsuko Sekimori, Risa Fujinaga, and Erina Ishikawa for technical assistance of sequence. Mika Kobayashi, Tomoko Tanaka, and Chie Akaishi for assistance. A.I. was a doctoral student fellow of Fostering Advanced Human Resources to Lead Green Transformation (GX)(SPRING GX) in the University of Tokyo. This work was supported by Core Research for Evolutional Science and Technology (CREST) of the Japan Science and Technology Agency (JST), Japan (JPMJCR13W2), Grant-in-Aid for the Japan Society for the Promotion of Science (JSPS) KAKENHI grant numbers 19H01048, 21K19519, 22H04991 (to H.K.), 19K08308, 22K07877 (to S. M.-T.), 22K20917, 24K18996 (to A.I.) 24K11186 (to Y.K.), 17J11177, 20H04858, 20K15858 (to H.H.), 16H06279 (PAGS), JP22H04925 (PAGS), the Mitsubishi foundation, the Fugaku Foundation (to H.K.) and RIKAKEN HD Life science research grant (to A.I.).

## Author contributions

A.I., Y.U., D.S., Y.K., S.M.-T., and H.Kurihara. conceived the study and designed experiments. A.I., Y.U., D.S., M.K., H.H., K.M., A.T, Y.H., and Y.W. performed experiments. C.S., Y.U., T. Kawamura, O.N., H.Katoh, S.I. and Y.K. supported the experiments. A.T., S.Y., S.F., T.K., M.S., A. K., Y.S., Y.W., and H. Aburatani provided the sequence platform. A.I., Y.U., S.Y., S.F., T.Kohro., and S.N. analyzed the sequence data. C.S. and H. Akiyama provided mutant mice. A.I. and H.Kurihara. wrote the manuscript with help from other authors.

## Materials & Correspondence

A.I. and H.K. are responsible for materials and correspondence related to this study. C.S. and H.Akiyama provided the S*ox9EGPP, ScxTomato, Sox9CreERT2, ScxCreL, ScxCreERT2,* and *Ai14* mutant mice.

## Supplemental figure legends

**Figure S1 Transcriptomic and chromatin accessibility profiles of diverse cell types in the cardiopharyngeal region.**

(a) FACS gating of single-cell suspension from *Wnt1-Cre; R26R-EYFP* embryos. All events were hierarchical gated by forward scatter (FSC) and side scatter (SSC) including area (A), height (H), and width (W). FSC singlets were gated, followed by exclusion of 7-AAD-positive dead cells to sort live EYFP-positive NCCs (P3) and EYFP-negative non-NCCs (P2).

(b) Heatmap showing differentially expressed genes (DEGs) across cardiopharyngeal cell clusters (as in Fig. 1d). DEGs with an average log2 expression > 1 were ranked by adjusted p-value from the Wilcoxon rank-sum test with Bonferroni correction, and the top ten genes per cluster were selected. Values represent cluster-averaged expression.

(c, d) Heatmaps showing representative TFs differentially expressed in each cluster (c) and the enrichment of their binding motifs in open chromatin regions (d) across cardiopharyngeal cell clusters.

(e, f) Heatmaps comparing differentially expressed TF genes (e) and the enrichment of their binding motifs in open chromatin regions (f) between NCC-derived and non-NCC-derived mesenchymal cells at E14.5 and E17.5.

**Figure S2 Xenium analysis of cardiopharyngeal tissues reveals distinct transcriptomic signatures across diverse cell types.**

(a-i) Xenium spatial transcriptomics of cardiopharyngeal tissues at E11.5 (a, b) and E12.5 (c-i) with hematoxylin-eosin staining.

(j) Heatmaps showing DEGs across Xenium-identified cell clusters (as in Fig. 2e).

(k, l) β-Galactosidase staining of cardiopharyngeal tissues from *Wnt1-Cre;R26R-LacZ* embryos at E11.5 (k) and E12.5 (l).

(m-p) Comparison of NCC-derived (m, n) and non-NCC-derived (o, p) mesenchymal cell distributions between fine cluster identities (as in Figure 1d) and RCTD-predicted cell clusters at E11.5 (m, n) and E12.5 (o, p).

(q, s) Estimated mean *EYFP* expression by Tangram across Xenium integrated clusters. The horizontal lines indicate the threshold corresponding to the 65th percentile across all spots.

**Figure S3 Transcriptional and developmental profiling of NCC-derived lineages.**

(a) Heatmap showing DEGs across integrated NCC clusters (as in Fig. 3b). DEGs with an average log2 expression > 1 were ranked by adjusted p-value from the Wilcoxon rank-sum test with Bonferroni correction, and the top ten genes per cluster were selected. Values represent cluster-averaged expression.

(b) UMAP decomposition of NCC subcategories (as in Fig. 4a) across sequential developmental stages.

(c) Cell type classification at each developmental stage.

(d) Cell cycle phase classification of each cluster.

(e-l) Feature plots showing representative marker gene expression for distinct NCC populations; neural tube (premigratory) (e), neural (f), glial/melanocyte (g), pharyngeal mesenchyme (h), intracardiac mesenchyme (i), smooth muscle (great arteries and ductus arteriosus) (j), and coronary artery smooth muscle (k).

(l) Immunostaining for Myh11 (red) and Dlk1 (blue). NCCs are marked by EYFP (green) and in *Wnt1-Cre;R26R* embryos at E17.5. Nuclei are stained by DAPI (white). Boxed regions (top) are shown at higher magnification (bottom). Scale bars, 50μm.

**Figure S4 Characterization of NCC-derived smooth muscle cell (SMC) subpopulations.**

(a) UMAP plot colored corresponding SMC subpopulation.

(b-d) Feature plots showing representative marker gene expression for distinct SMC populations; *Myh11* (mature SMCs) (b), *Sost* (great artery SMCs) (c), and *Reln* (coronary artery SMCs) (d).

(f) Violin plot of Myh11 across SMC subclusters.

(g) Heatmap showing DEGs across SMC subpopulations. DEGs with an average log2 expression > 2.5 were ranked by adjusted p-value from the Wilcoxon rank-sum test with Bonferroni correction, and the top 30 genes per cluster were selected. Values represent cluster-averaged expression.

(h-s) Immunostaining for Myh11 (green) (h, k, n, q) with Sost (red) (i, j, l, m) or Reln (red) (o, r, p, s). Nuclei are stained by DAPI (blue) at E17.5 embryos. Ao, aorta; DA; dorsal aorta; RCA. right coronary artery; SB, septal branch. Scale bars, 50μm.

**Figure S5 Characterization of NCC-derived intracardiac mesenchyme.**

(a) UMAP plot colored corresponding intracardiac mesenchymal subpopulation.

(b, c) Heatmaps showing DEGs across intracardiac mesenchymal subpopulations for genes included (b) or not included (c) in the customized Xenium panel. DEGs with an average log2 expression > 0.15 were ranked by adjusted p-value from the Wilcoxon rank-sum test with Bonferroni correction, and the top 25 genes per cluster were selected. Values represent cluster-averaged expression.

(d, e) Feature plots showing the expression of *Tcf24* (d) and *Vegfc* (e).

(f-j) Xenium spatial transcriptomic visualization of *Tcf24*, *Vegfc*, and *Postn* (g-j) with corresponding hematoxylin-eosin staining (f) in E12.5 cardiac tissue. Ao, aorta; AP septum, aorticopulmonary septum; PA, pulmonary artery; RA, right atrium. Scale bars, 500 μm.

**Figure S6 Visium spatial transcriptomics of the *Wnt1-Cre*-labeled developing heart.**

Spatial transcriptomic profiling using the 10x Visium platform at E14.5 (a-f) and E17.5 (g-l). On hematoxylin-eosin-stained images (a, g), clustering profiles are visualized (b, h) with UMAP plots, colored corresponding to early Visium clusters (evCs) (c), late Visium clusters (lvCs) (i) and stages (d, j). EYFP-expressing spots are overlaid on the tissue sections (e, k), with corresponding cluster annotations (f, l).

**Figure S7 Spatial localization of NCC-derived mesenchymal clusters on the Visium datasets.**

Spatial mapping of NCC-derived mesenchymal clusters on the Visium datasets at E14.5 (a, b) and E17.5 (c, d), highlighting the predicted correspondence between clusters and spatial spots. Color bars represent the probability of spatial prediction.

**Figure S8 *Hox* gene expression patterns in NCC-derived lineages.**

Feature plots showing the expression of *Hox* genes in NCC-derived lineages.

**Figure S9 Decoding *Hox* codes in NCC-derived lineages and dynamics of gene expression profiles at bifurcation point.**

(a) Schematic illustration of *Hox*-code decoding during development. Cells expressing any *Hox2* paralog, but lacking *Hox3–5* paralogs, were defined as PA2-derived preotic NCCs, whereas cells expressing any of *Hox3*–5 paralogs were classified as PA3/4/6-derived postotic NCCs. Preotic, postotic, and *Hox*-negative populations were then projected onto the integrated UMAP across developmental stages (E10.5–E14.5). Three flows of NCC contribution to the cardiovascular system were inferred across developmental stages.

(b–f) Trajectory inference based on the Slingshot (b, e) and scVelo (c, f) and heatmaps showing gene expression dynamics (d, g) of NCC-derived lineages at E10.5 and E11.5 (b–d) or E12.5, E13.5, and E14.5 (e, f, g). Trajectory was set from C14 to each lineage, and heatmaps showed each top 30 DEGs according to lineages (d, g). DEGs were calculated as Wald statistics accompanied with pseudotime and filtered with an average log2 expression > 1, and the top 30 genes with the result of Wald statistics per cluster were selected. Color scales indicate z-scored values for each gene.

**Figure S10 Characterization of NCC-derived mesenchymal clusters by GRNs represented as overall communities (OCs).**

(a) Projection of Fluidigm C1 scRNA-seq data onto the integrated UMAP, colored by subcategorization of NCC clusters as shown in Fig. 4a.

(b) Heatmap showing the enrichment of OCs across integrated mesenchymal clusters. Color scale represents z-scored enrichment values.

(c) Key OCs regulating NCC lineage commitment, identified using a random forest classifier.

(d) Confusion matrix showing the performance of the random forest-based model in predicting NCC cluster identity.

**Figure S11 Chromatin accessibility of *Hoxa2* super-enhancer in NCC-derived mesenchymal clusters**

(a-c) Coverage plots of chromatin accessibility at the *Hoxa2* super-enhancer loci (*Hoxa* inter-TAD regulatory element (HIRE) 1 and HIRE2) located 1.33 Mb away from the *Hoxa2* locus in NCC-derived mesenchymal subpopulations. Magnified views of HIRE1(chr6:50,913,170-51,087,888) and HIRE2 (chr6:50,789,172-50,828,639) are highlighted in gray (b, c). Differentially accessible peaks (DAPs) are highlighted in orange (pharyngeal and transitional mesenchyme) and blue (cardiac cushion and subvalvular mesenchyme) (b). Transcription factor-binding motifs are indicated in pink (Meis1; MA1639.2) and green (Gata4; MA0482.3). Bigwig tracks of bulk ATAC-seq signals in the mandibular arch (Md) and second pharyngeal arch (PA2) at E10.5. are shown.

(d, e) Top 12 enriched TF-binding motifs identified from DAPs within HIRE1 and HIRE2 in pharyngeal and transitional mesenchyme (d) and cardiac cushion and subvalvular mesenchyme (e).

**Figure S12 Histone modifications and chromatin accessibility at the *Sox9* locus.**

(a) ChIP-seq and ATAC-seq data of the E10.5 to E116.5 mouse embryonic heart from ENCODE, accessed via the UCSC Genome Browser (http://genome.ucsc.edu/). The candidate distal enhancer region is highlighted in orange.

(b) Coverage plots of chromatin accessibility at *Sox9* loci in cardiopharyngeal cell clusters in Fig. 1e (left), with violin plots showing gene expression from scRNA-seq data (upper right). The candidate distal enhancer region is highlighted in light green. Note that Sox9 expression and accessibility to the enhancer region are upregulated in *Tbx20*^high^ mesenchymal (cluster 3) and epicardial (cluster 5) cells.

**Figure S13 *Sox9* expression and open chromatin peaks in *Wt1*^high^ and *Wt1*^low^ cell population.**

(a) UMAP plot of *Wt*1^high^ (C22) and *Wt1*^low^ (C5) clusters extracted from Figure 1d.

(b, c) Feature plots showing the expression of *Sox9* (b) and *Wt1* (c).

(d) Coverage plot of chromatin accessibility at the *Sox9* locus in *Wt1*^high^ and *Wt1*^low^ cell populations (left), with violin plots showing *Sox9* expression from scRNA-seq data (right).

## References

1. Martik, M. L. & Bronner, M. E. Riding the crest to get a head: neural crest evolution in vertebrates. Nat. Rev. Neurosci. 22, 616–626 (2021).

2. Etchevers, H. C., Dupin, E. & Le Douarin, N. M. The diverse neural crest: from embryology to human pathology. Development 146, dev169821 (2019).

3. Le Douarin, N. & Kalcheim, C. The Neural Crest. (Cambridge University Press, Cambridge, 1999). doi:10.1017/CBO9780511897948.

4. Kirby, M. L., Gale, T. F. & Stewart, D. E. Neural Crest Cells Contribute to Normal Aorticopulmonary Septation. Science (1979). 220, 1059–1061 (1983).

5. Kirby, M. L. & Hutson, M. R. Factors controlling cardiac neural crest cell migration. Cell Adh. Migr. 4, 609–621 (2010).

6. Waldo, K., Miyagawa-Tomita, S., Kumiski, D. & Kirby, M. L. Cardiac Neural Crest Cells Provide New Insight into Septation of the Cardiac Outflow Tract: Aortic Sac to Ventricular Septal Closure. Dev. Biol. 196, 129–144 (1998).

7. Nishibatake, M., Kirby, M. L. & Van Mierop, L. H. Pathogenesis of persistent truncus arteriosus and dextroposed aorta in the chick embryo after neural crest ablation. Circulation 75, 255–264 (1987).

8. Arima, Y. et al. Preotic neural crest cells contribute to coronary artery smooth muscle involving endothelin signalling. Nat. Commun. 3, 1267 (2012).

9. Miyagawa-Tomita, S., Arima, Y. & Kurihara, H. The “Cardiac Neural Crest” Concept Revisited. in Etiology and Morphogenesis of Congenital Heart Disease 227–232 (Springer Japan, Tokyo, 2016). doi:10.1007/978-4-431-54628-3_30.

10. Soldatov, R. et al. Spatiotemporal structure of cell fate decisions in murine neural crest. Science (1979). 364, eaas9536 (2019).

11. Rothstein, M., Bhattacharya, D. & Simoes-Costa, M. The molecular basis of neural crest axial identity. Dev. Biol. 444, S170–S180 (2018).

12. Gandhi, S., Ezin, M. & Bronner, M. E. Reprogramming Axial Level Identity to Rescue Neural-Crest-Related Congenital Heart Defects. Dev. Cell 53, 300–315.e4 (2020).

13. De Bono, C. et al. Single-cell transcriptomics uncovers a non-autonomous Tbx1-dependent genetic program controlling cardiac neural crest cell development. Nat. Commun. 14, (2023).

14. Chen, W. et al. Single□cell transcriptomic landscape of cardiac neural crest cell derivatives during development. EMBO Rep. 22, 1–18 (2021).

15. Tomita, Y. et al. Cardiac neural crest cells contribute to the dormant multipotent stem cell in the mammalian heart. J. Cell Biol. 170, 1135–1146 (2005).

16. Tamura, Y. et al. Neural Crest–Derived Stem Cells Migrate and Differentiate Into Cardiomyocytes After Myocardial Infarction. Arterioscler. Thromb. Vasc. Biol. 31, 582–589 (2011).

17. Biancalani, T. et al. Deep learning and alignment of spatially resolved single-cell transcriptomes with Tangram. Nat. Methods 18, 1352–1362 (2021).

18. Jiang, X., Rowitch, D. H., Soriano, P., McMahon, A. P. & Sucov, H. M. Fate of the mammalian cardiac neural crest. Development 127, 1607–1616 (2000).

19. Cable, D. M. et al. Robust decomposition of cell type mixtures in spatial transcriptomics. Nat. Biotechnol. 40, 517–526 (2022).

20. Fu, M. et al. Neural Crest Cells Differentiate Into Brown Adipocytes and Contribute to Periaortic Arch Adipose Tissue Formation. Arterioscler. Thromb. Vasc. Biol. 39, 1629–1644 (2019).

21. Zhao, F., Bosserhoff, A.-K., Buettner, R. & Moser, M. A Heart-Hand Syndrome Gene: Tfap2b Plays a Critical Role in the Development and Remodeling of Mouse Ductus Arteriosus and Limb Patterning. PLoS One 6, e22908 (2011).

22. Yokoyama, U. et al. Chronic activation of the prostaglandin receptor EP4 promotes hyaluronan-mediated neointimal formation in the ductus arteriosus. Journal of Clinical Investigation 116, 3026–3034 (2006).

23. Volz, K. S. et al. Pericytes are progenitors for coronary artery smooth muscle. Elife 4, 1–22 (2015).

24. Webb, S., Qayyum, S. R., Anderson, R. H., Lamers, W. H. & K. Richardson, M. Septation and separation within the outflow tract of the developing heart. J. Anat. 202, 327–342 (2003).

25. Sizarov, A. et al. Three□dimensional and molecular analysis of the arterial pole of the developing human heart. J. Anat. 220, 336–349 (2012).

26. Anderson, R. H., Mori, S., Spicer, D. E., Brown, N. A. & Mohun, T. J. Development and Morphology of the Ventricular Outflow Tracts. World J. Pediatr. Congenit. Heart Surg. 7, 561–577 (2016).

27. Minoux, M. & Rijli, F. M. Molecular mechanisms of cranial neural crest cell migration and patterning in craniofacial development. Development 137, 2605–2621 (2010).

28. Parker, H. J., Pushel, I. & Krumlauf, R. Coupling the roles of Hox genes to regulatory networks patterning cranial neural crest. Dev. Biol. 444, S67–S78 (2018).

29. Street, K. et al. Slingshot: cell lineage and pseudotime inference for single-cell transcriptomics. BMC Genomics 19, 477 (2018).

30. Weiler, P., Lange, M., Klein, M., Pe’er, D. & Theis, F. CellRank 2: unified fate mapping in multiview single-cell data. Nat. Methods 21, 1196–1205 (2024).

31. Bobola, N. & Sagerström, C. G. TALE transcription factors: Cofactors no more. Semin. Cell Dev. Biol. 152–153, 76–84 (2024).

32. Bridoux, L. et al. HOX paralogs selectively convert binding of ubiquitous transcription factors into tissue-specific patterns of enhancer activation. PLoS Genet. 16, e1009162 (2020).

33. Penkov, D. et al. Analysis of the DNA-Binding Profile and Function of TALE Homeoproteins Reveals Their Specialization and Specific Interactions with Hox Genes/Proteins. Cell Rep. 3, 1321–1333 (2013).

34. Tamada, Y. et al. Estimating Genome-Wide Gene Networks Using Nonparametric Bayesian Network Models on Massively Parallel Computers. IEEE/ACM Trans. Comput. Biol. Bioinform. 8, 683–697 (2011).

35. Aibar, S. et al. SCENIC: single-cell regulatory network inference and clustering. Nat. Methods 14, 1083–1086 (2017).

36. Kalinka, A. T. & Tomancak, P. linkcomm: an R package for the generation, visualization, and analysis of link communities in networks of arbitrary size and type. Bioinformatics 27, 2011–2012 (2011).

37. Kessler, S. et al. A multiple super-enhancer region establishes inter-TAD interactions and controls Hoxa function in cranial neural crest. Nat. Commun. 14, 3242 (2023).

38. Machon, O., Masek, J., Machonova, O., Krauss, S. & Kozmik, Z. Meis2 is essential for cranial and cardiac neural crest development. BMC Dev. Biol. 15, 40 (2015).

39. Akiyama, H. et al. Essential role of Sox9 in the pathway that controls formation of cardiac valves and septa. Proceedings of the National Academy of Sciences 101, 6502–6507 (2004).

40. Liu, H. et al. Odd-skipped related-1 controls neural crest chondrogenesis during tongue development. Proceedings of the National Academy of Sciences 110, 18555–18560 (2013).

41. Kamimoto, K. et al. Dissecting cell identity via network inference and in silico gene perturbation. Nature 614, 742–751 (2023).

42. Roux, M. et al. Hoxa1 and Hoxb1 are required for pharyngeal arch artery development. Mech. Dev. 143, 1–8 (2017).

43. Zaffran, S., Odelin, G., Stefanovic, S., Lescroart, F. & Etchevers, H. C. Ectopic expression of *Hoxb1* induces cardiac and craniofacial malformations. genesis 56, (2018).

44. Chisaka, O. & Capecchi, M. R. Regionally restricted developmental defects resulting from targeted disruption of the mouse homeobox gene hox-1.5. Nature 350, 473–479 (1991).

45. Kameda, Y., Watari-Goshima, N., Nishimaki, T. & Chisaka, O. Disruption of the Hoxa3 homeobox gene results in anomalies of the carotid artery system and the arterial baroreceptors. Cell Tissue Res. 311, 343–352 (2003).

46. Mandel, E. M. et al. The BMP pathway acts to directly regulate Tbx20 in the developing heart. Development 137, 1919–1929 (2010).

47. Daoud, G. et al. BMP-mediated induction of GATA4/5/6 blocks somitic responsiveness to SHH. Development 141, 3978–3987 (2014).

48. Asai, R. et al. Amniogenic somatopleure: a novel origin of multiple cell lineages contributing to the cardiovascular system. Sci. Rep. 7, 8955 (2017).

49. Darieva, Z. et al. Ubiquitous MEIS transcription factors actuate lineage-specific transcription to establish cell fate. EMBO J. 44, 2232–2262 (2025).

50. Giffin, J. L., Gaitor, D. & Franz-Odendaal, T. A. The Forgotten Skeletogenic Condensations: A Comparison of Early Skeletal Development Amongst Vertebrates. J. Dev. Biol. 7, 4 (2019).

51. Hall, B. K. & Miyake, T. All for one and one for all: condensations and the initiation of skeletal development. BioEssays 22, 138–147 (2000).

52. Gao, Z. et al. Ets1 is required for proper migration and differentiation of the cardiac neural crest. Development 137, 1543–1551 (2010).

53. Arai, H. N. et al. Metalloprotease-Dependent Attenuation of BMP Signaling Restricts Cardiac Neural Crest Cell Fate. Cell Rep. 29, 603–616.e5 (2019).

54. Chen, D. et al. Fibronectin signals through integrin α5β1 to regulate cardiovascular development in a cell type-specific manner. Dev. Biol. 407, 195–210 (2015).

55. Sugimoto, Y., Takimoto, A., Hiraki, Y. & Shukunami, C. Generation and characterization of *ScxCre* transgenic mice. genesis 51, 275–283 (2013).

56. Blitz, E., Sharir, A., Akiyama, H. & Zelzer, E. Tendon-bone attachment unit is formed modularly by a distinct pool of Scx - and Sox9-positive progenitors. Development 140, 2680–2690 (2013).

57. Lincoln, J., Lange, A. W. & Yutzey, K. E. Hearts and bones: Shared regulatory mechanisms in heart valve, cartilage, tendon, and bone development. Dev. Biol. 294, 292–302 (2006).

58. Srinivas, S. et al. Cre reporter strains produced by targeted insertion of EYFP and ECFP into the ROSA26 locus. BMC Dev. Biol. 1, 4 (2001).

59. Furuyama, K. et al. Continuous cell supply from a Sox9-expressing progenitor zone in adult liver, exocrine pancreas and intestine. Nat. Genet. 43, 34–41 (2011).

60. Soeda, T. et al. Sox9-expressing precursors are the cellular origin of the cruciate ligament of the knee joint and the limb tendons. Genesis 48, 635–644 (2010).

61. Yu, X. et al. Dynamic interactions between cartilaginous and tendinous/ligamentous primordia during musculoskeletal integration. Development (Cambridge) 152, (2025).

62. Madisen, L. et al. A robust and high-throughput Cre reporting and characterization system for the whole mouse brain. Nat. Neurosci. 13, 133–140 (2010).

63. Harada, Y. et al. ETS-dependent enhancers for endothelial-specific expression of serum/glucocorticoid-regulated kinase 1 during mouse embryo development. Genes to Cells 26, 611–626 (2021).

64. Stuart, T., Srivastava, A., Madad, S., Lareau, C. A. & Satija, R. Single-cell chromatin state analysis with Signac. Nat. Methods 18, 1333–1341 (2021).

65. Hao, Y. et al. Integrated analysis of multimodal single-cell data. Cell 184, 3573–3587.e29 (2021).

66. Young, M. D. & Behjati, S. SoupX removes ambient RNA contamination from droplet-based single-cell RNA sequencing data. Gigascience 9, (2020).

67. McGinnis, C. S., Murrow, L. M. & Gartner, Z. J. DoubletFinder: Doublet Detection in Single-Cell RNA Sequencing Data Using Artificial Nearest Neighbors. Cell Syst. 8, 329–337.e4 (2019).

68. Schep, A. N., Wu, B., Buenrostro, J. D. & Greenleaf, W. J. ChromVAR: Inferring transcription-factor-associated accessibility from single-cell epigenomic data. Nat. Methods 14, 975–978 (2017).

69. Bergen, V., Lange, M., Peidli, S., Wolf, F. A. & Theis, F. J. Generalizing RNA velocity to transient cell states through dynamical modeling. Nat. Biotechnol. 38, 1408–1414 (2020).

70. La Manno, G. et al. RNA velocity of single cells. Nature 560, 494–498 (2018).

71. Schep, A. motifmatchr: Fast Motif Matching in R. Bioconductor version: Release (3. 12) Preprint at (2021).

72. Fornes, O. et al. JASPAR 2020: update of the open-access database of transcription factor binding profiles. Nucleic Acids Res. 48, D87–D92 (2019).

73. Siepel, A. et al. Evolutionarily conserved elements in vertebrate, insect, worm, and yeast genomes. Genome Res. 15, 1034–1050 (2005).

74. Ishii, M. et al. A Stable Cranial Neural Crest Cell Line from Mouse. Stem Cells Dev. 21, 3069–3080 (2012).

