## Supplementary figures and images for "*Hox*–*Meis*-relayed gene regulatory transition underlies cardiopharyngeal neural crest diversification"

### Supplemental figure 1

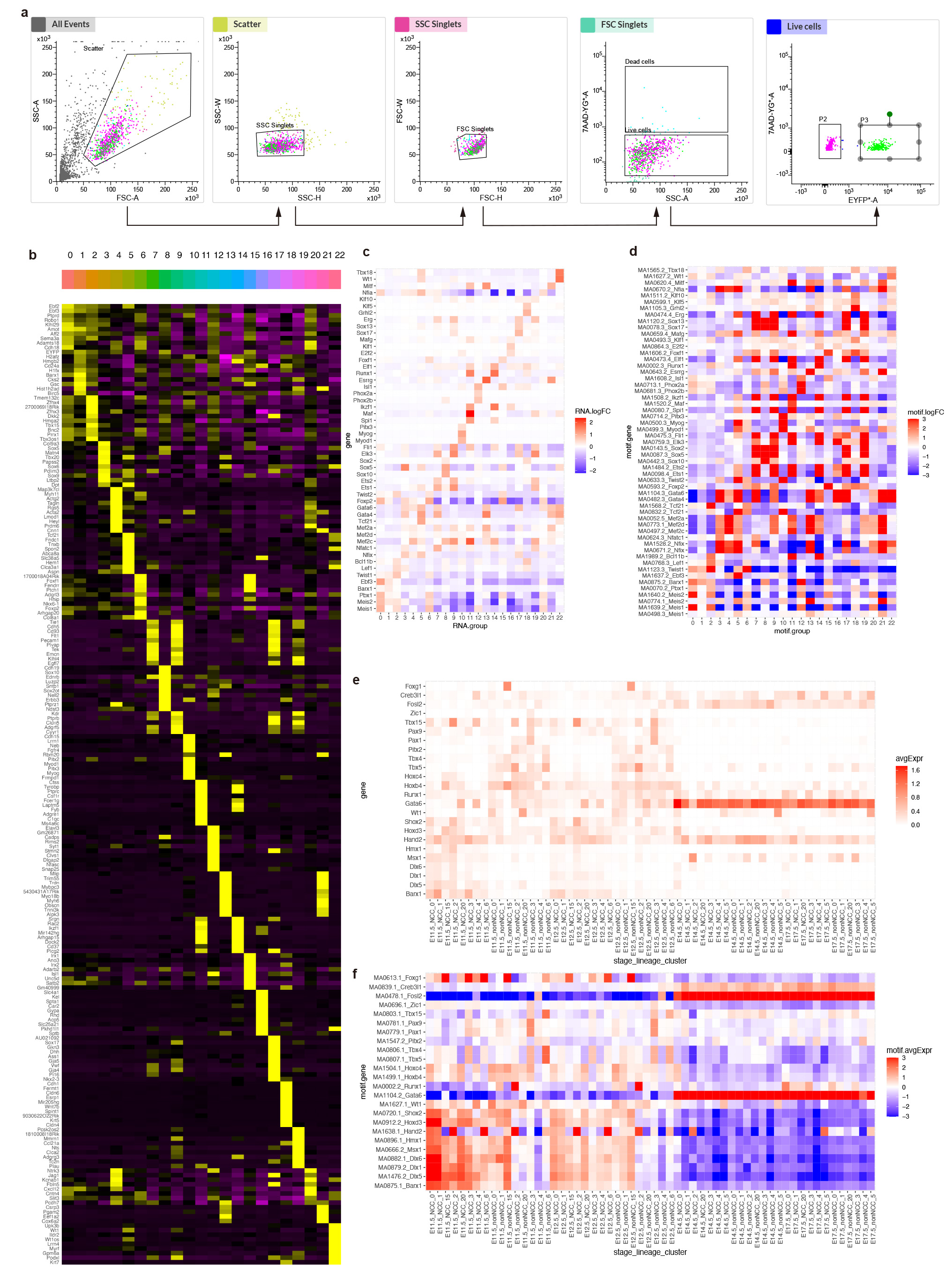

### Supplemental figure 2

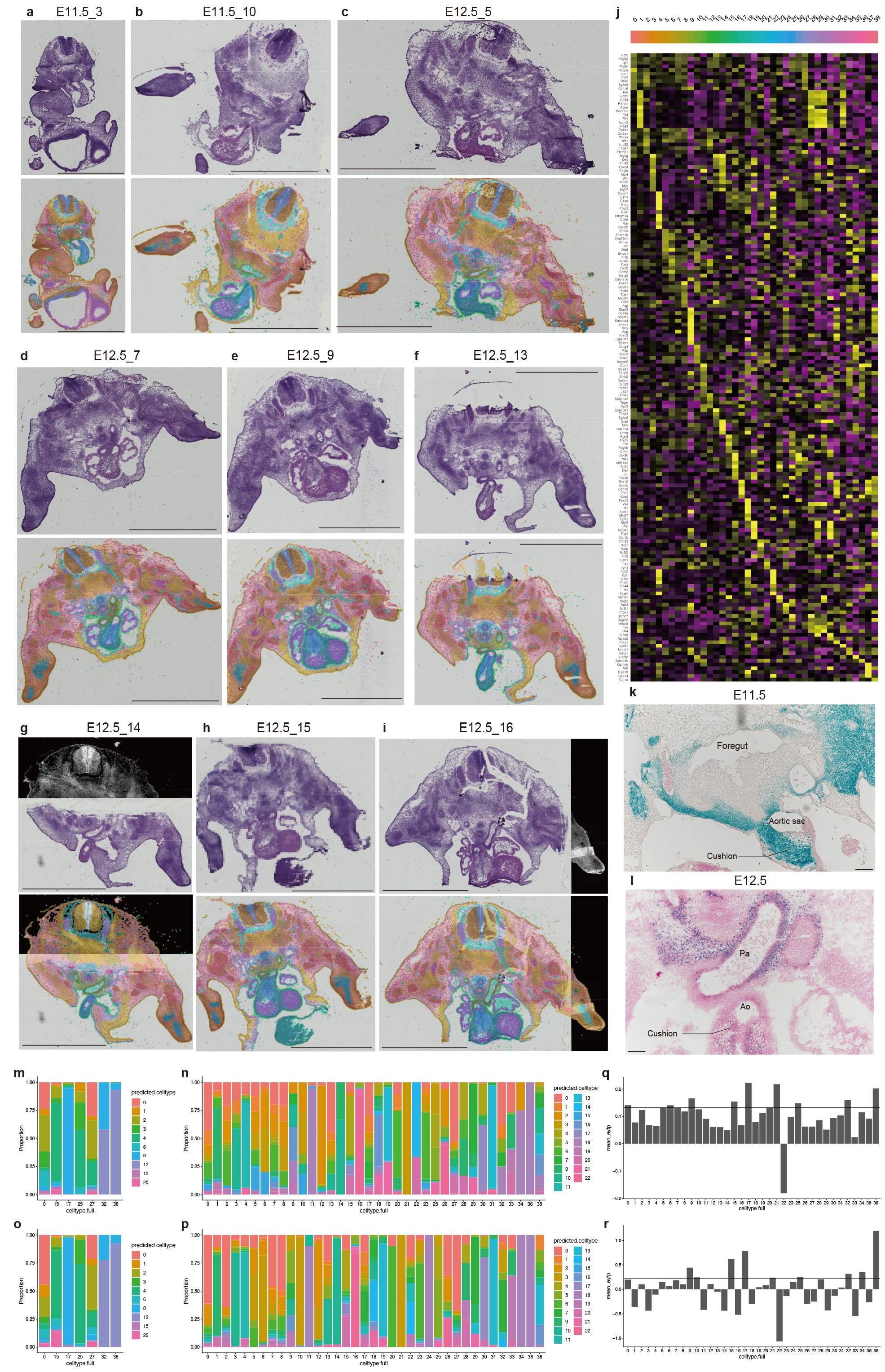

### Supplemental figure 3

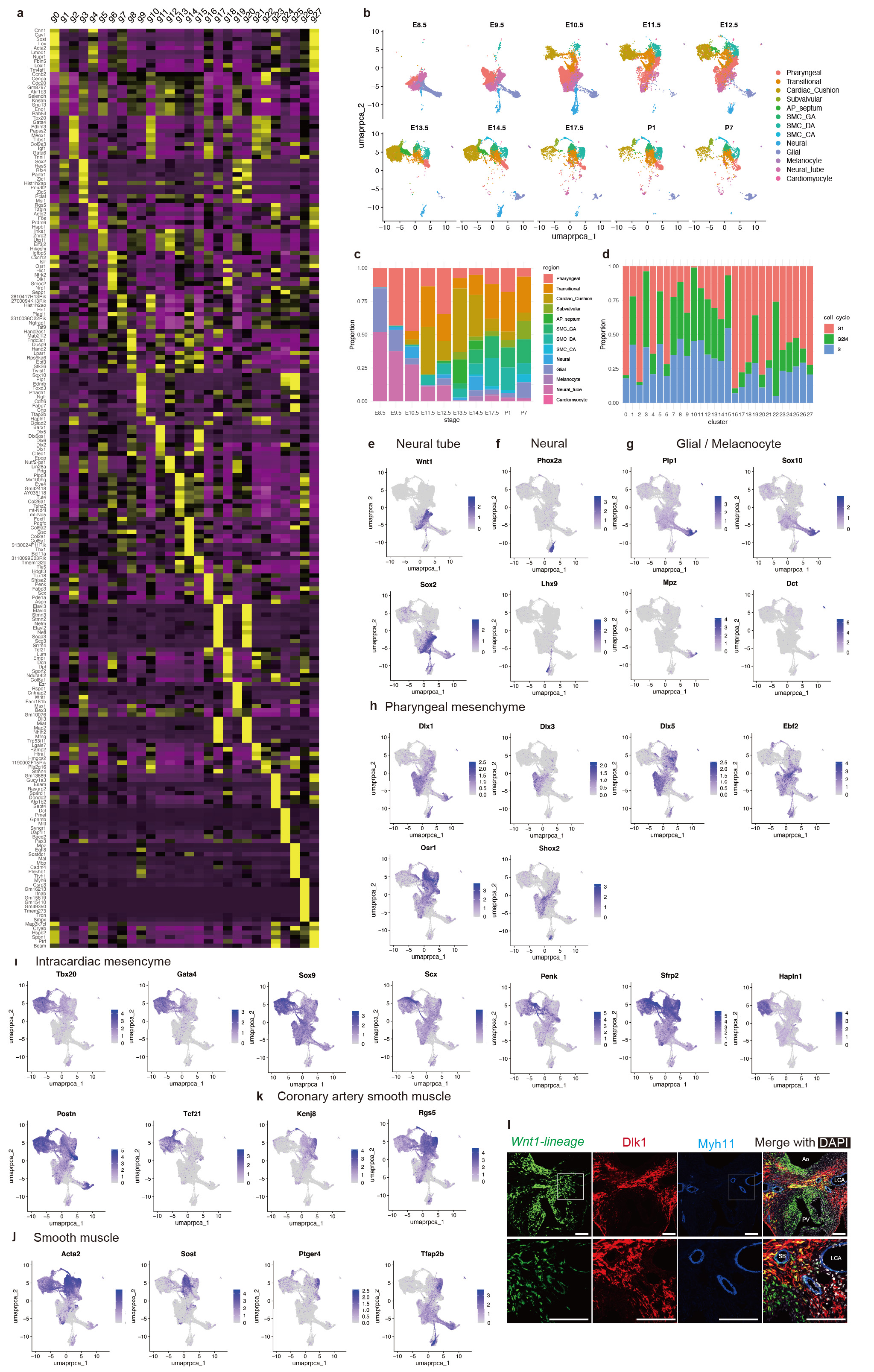

### Supplemental figure 4

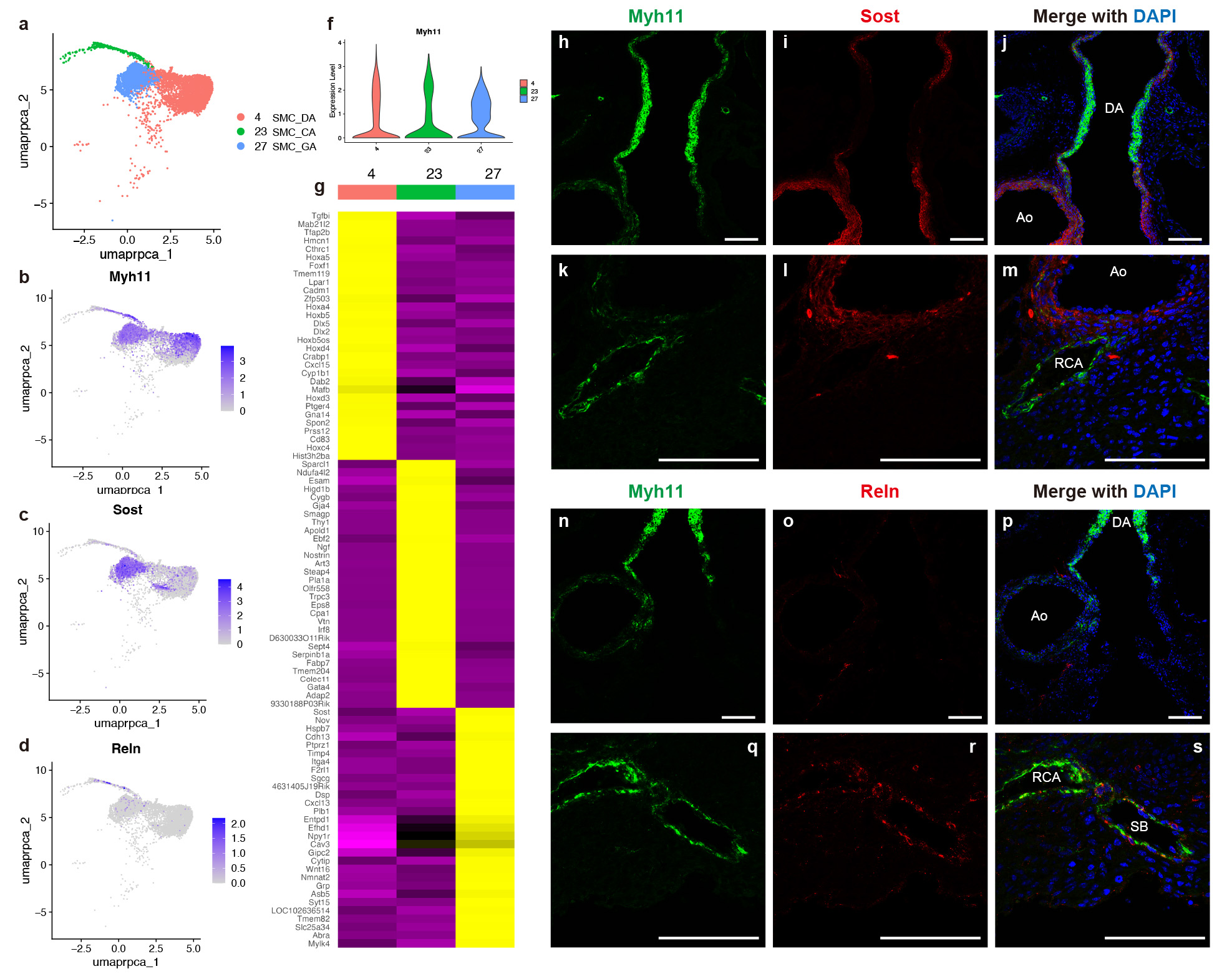

### Supplemental figure 5

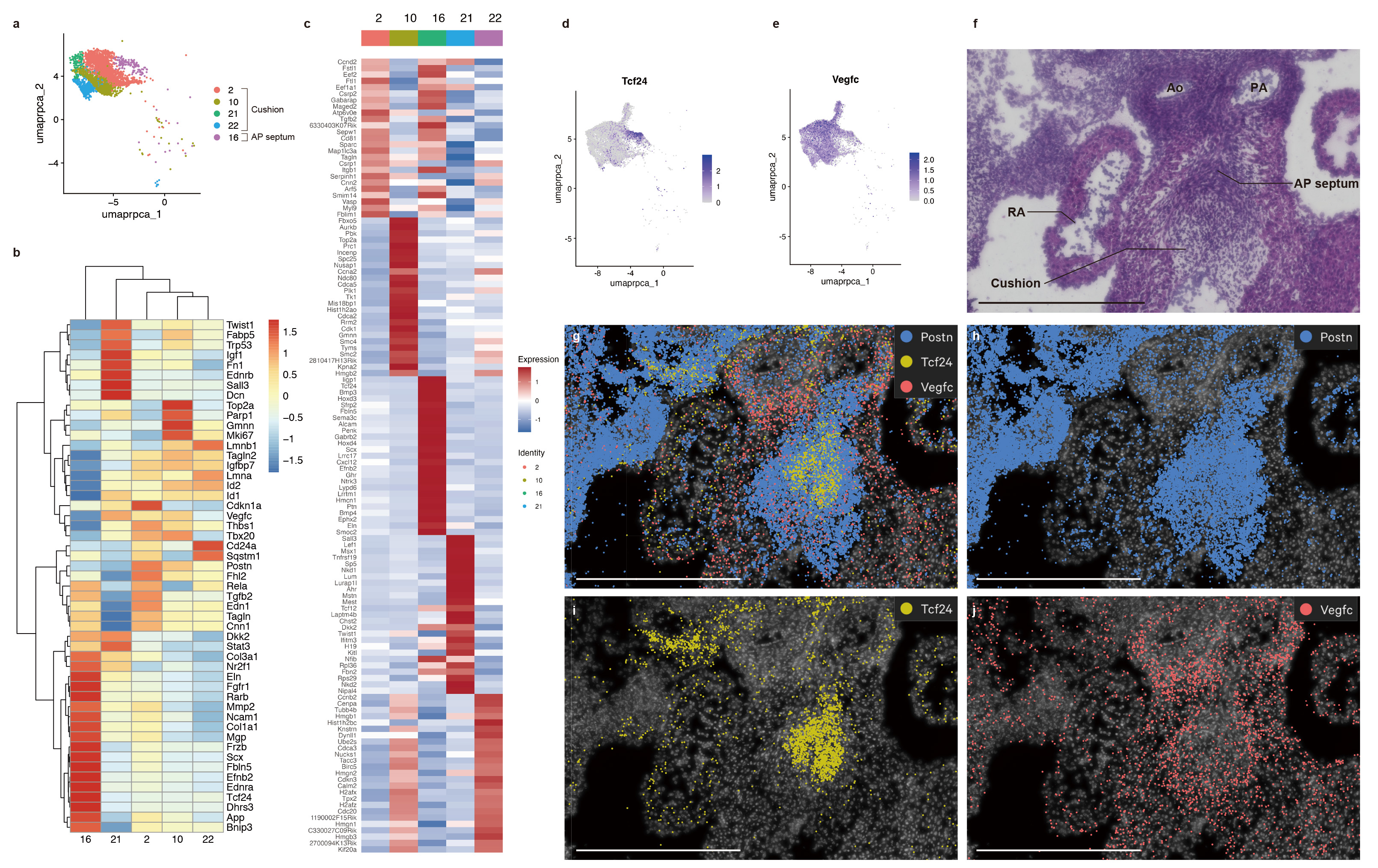

### Supplemental figure 6

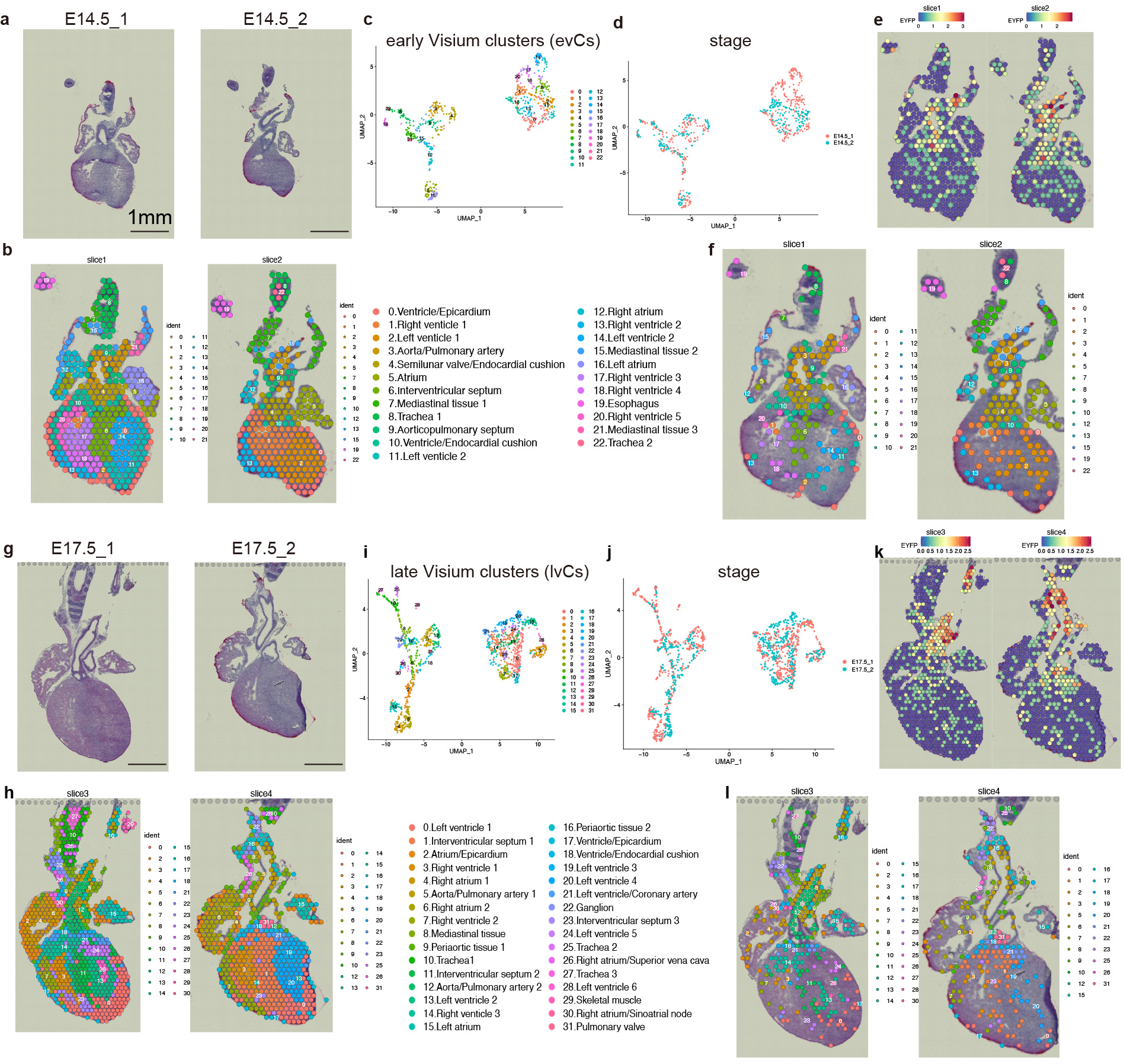

### Supplemental figure 7

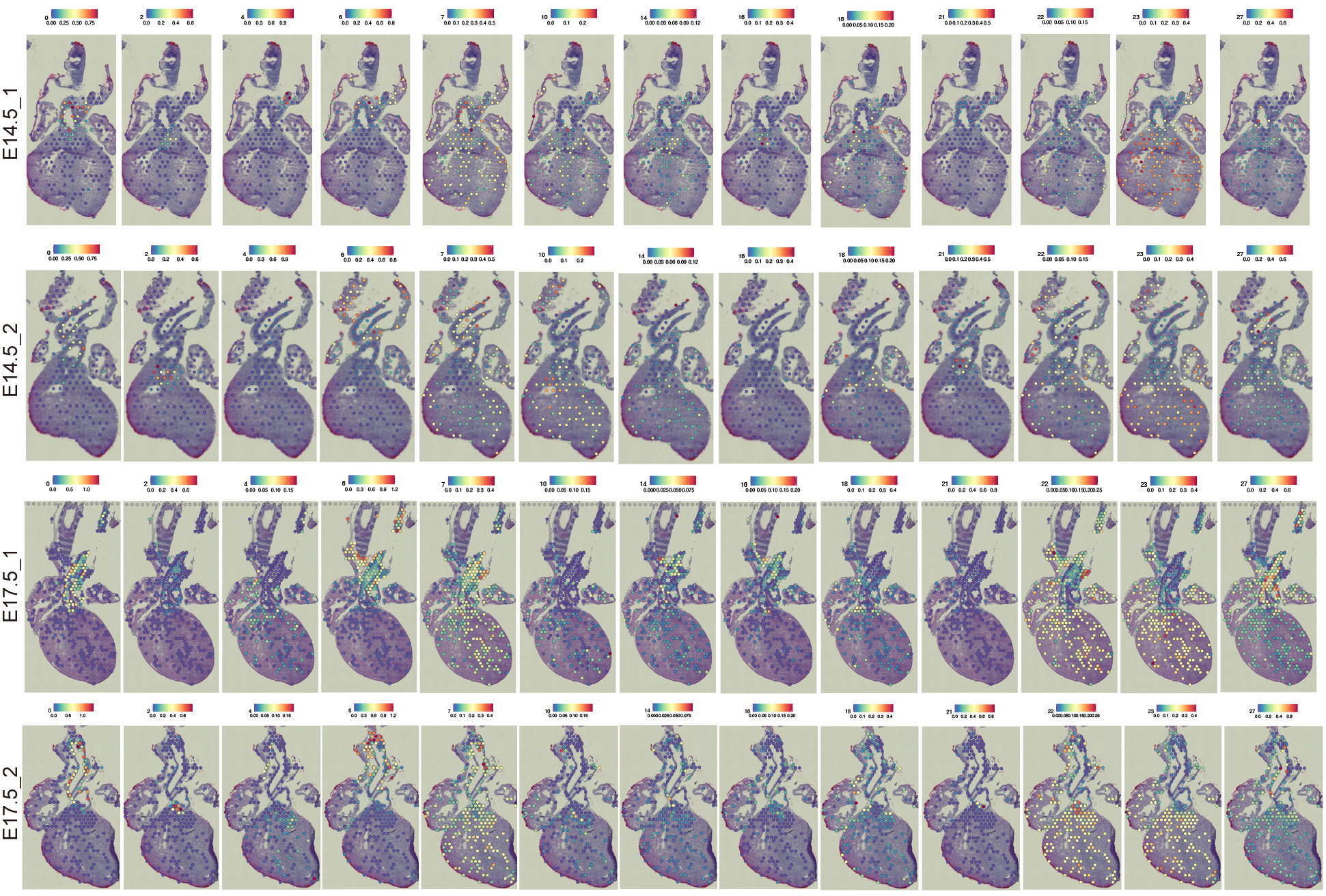

### Supplemental figure 8

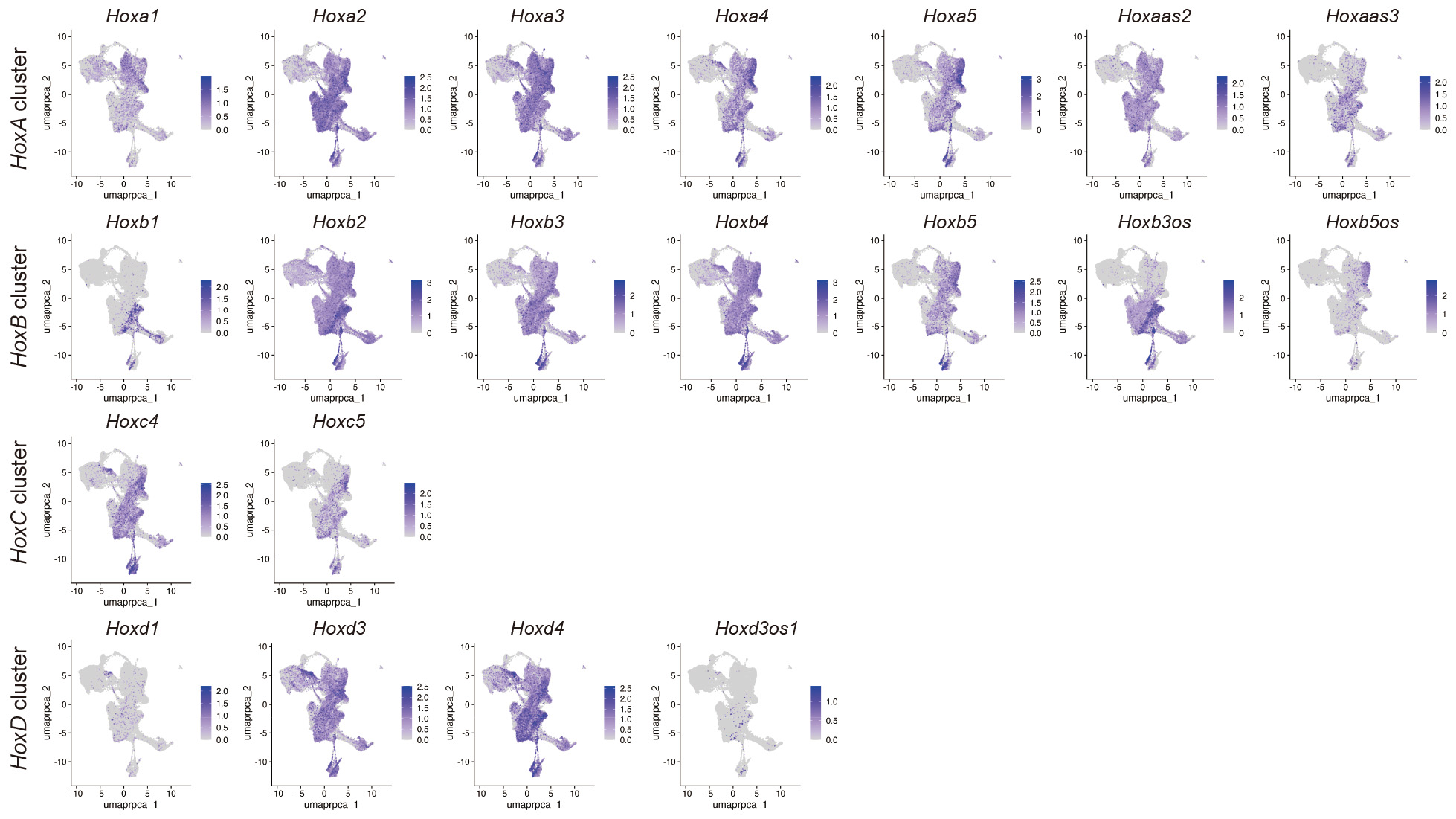

### Supplemental figure 9

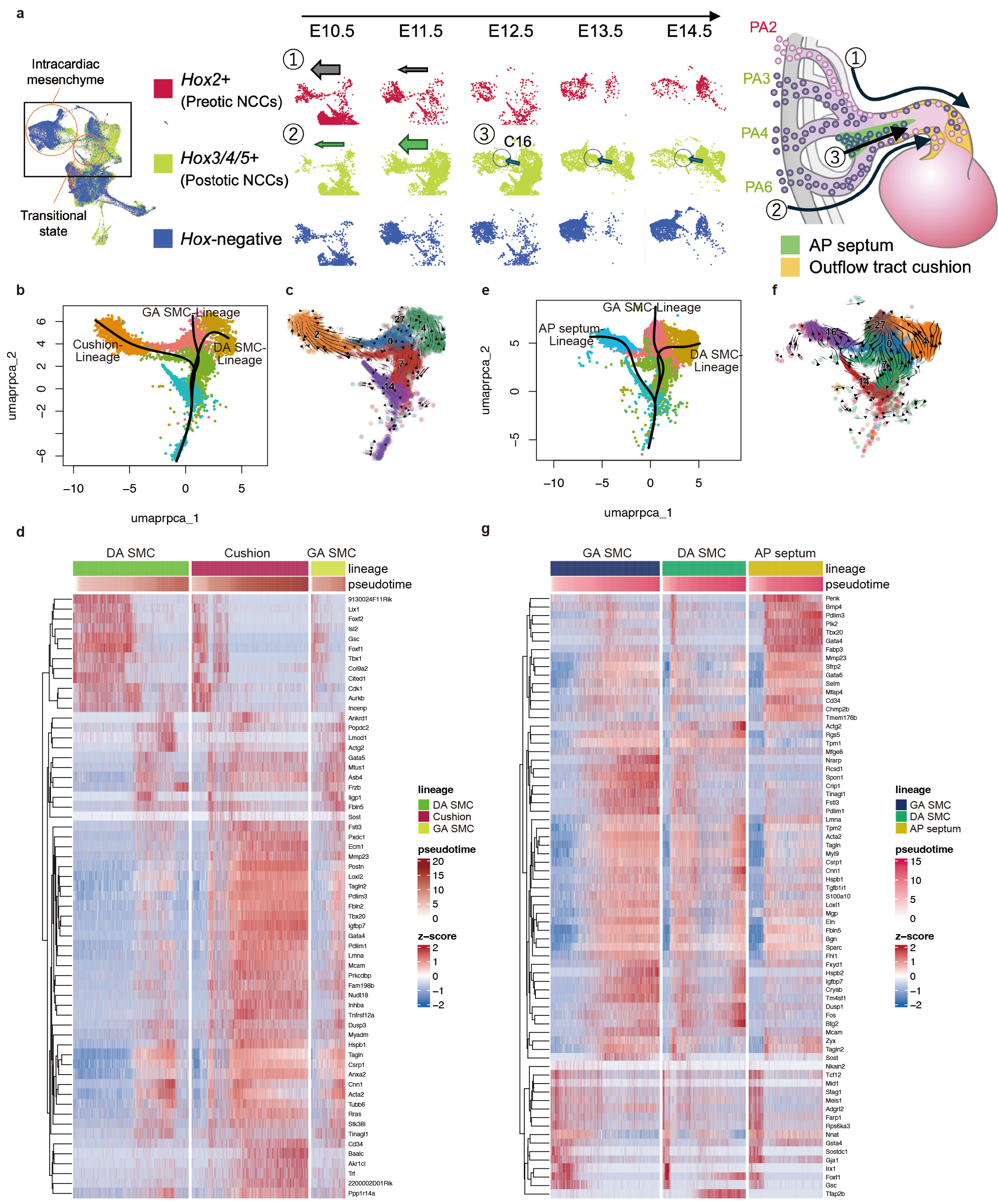

### Supplemental figure 10

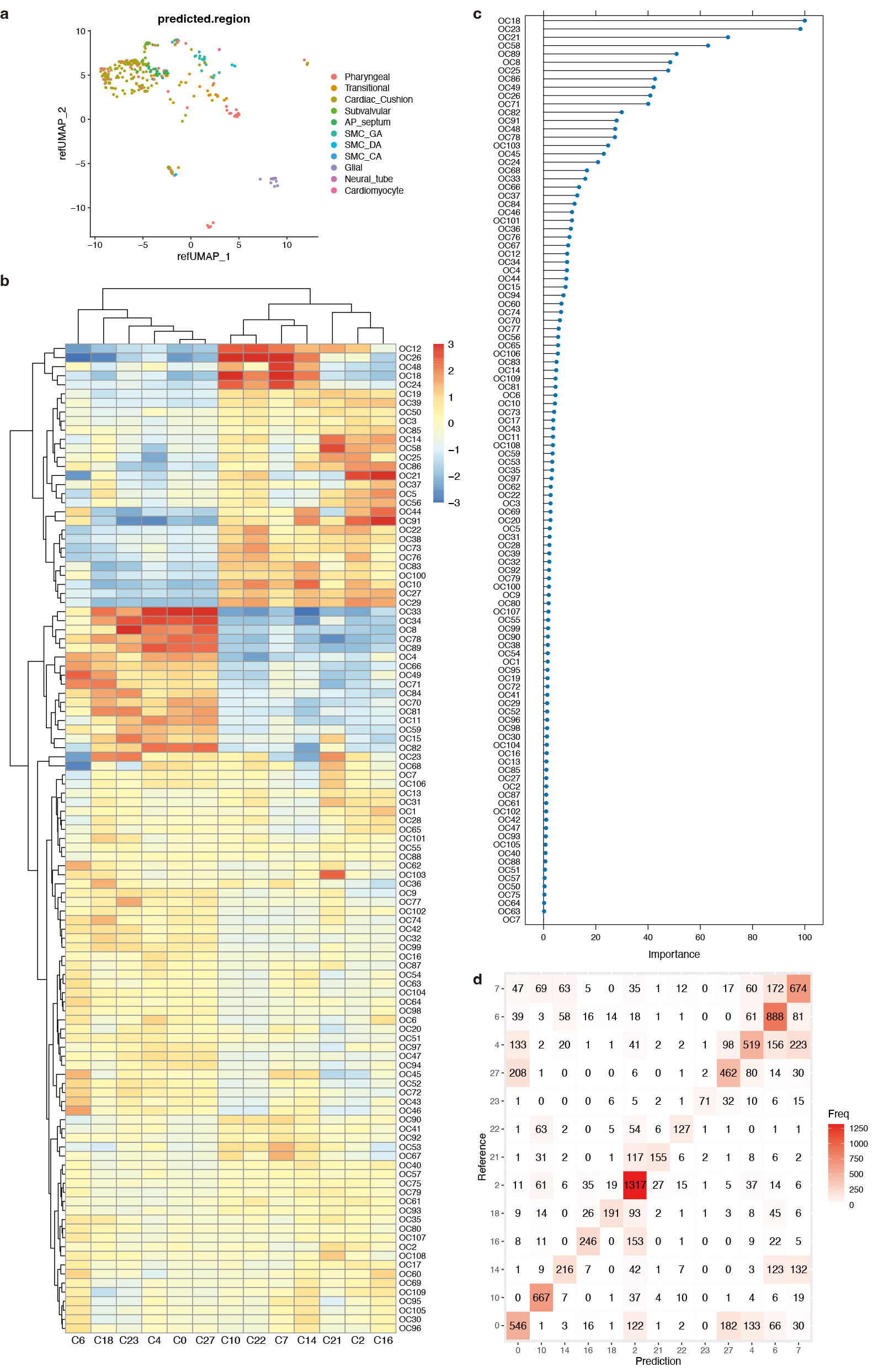

### Supplemental figure 11

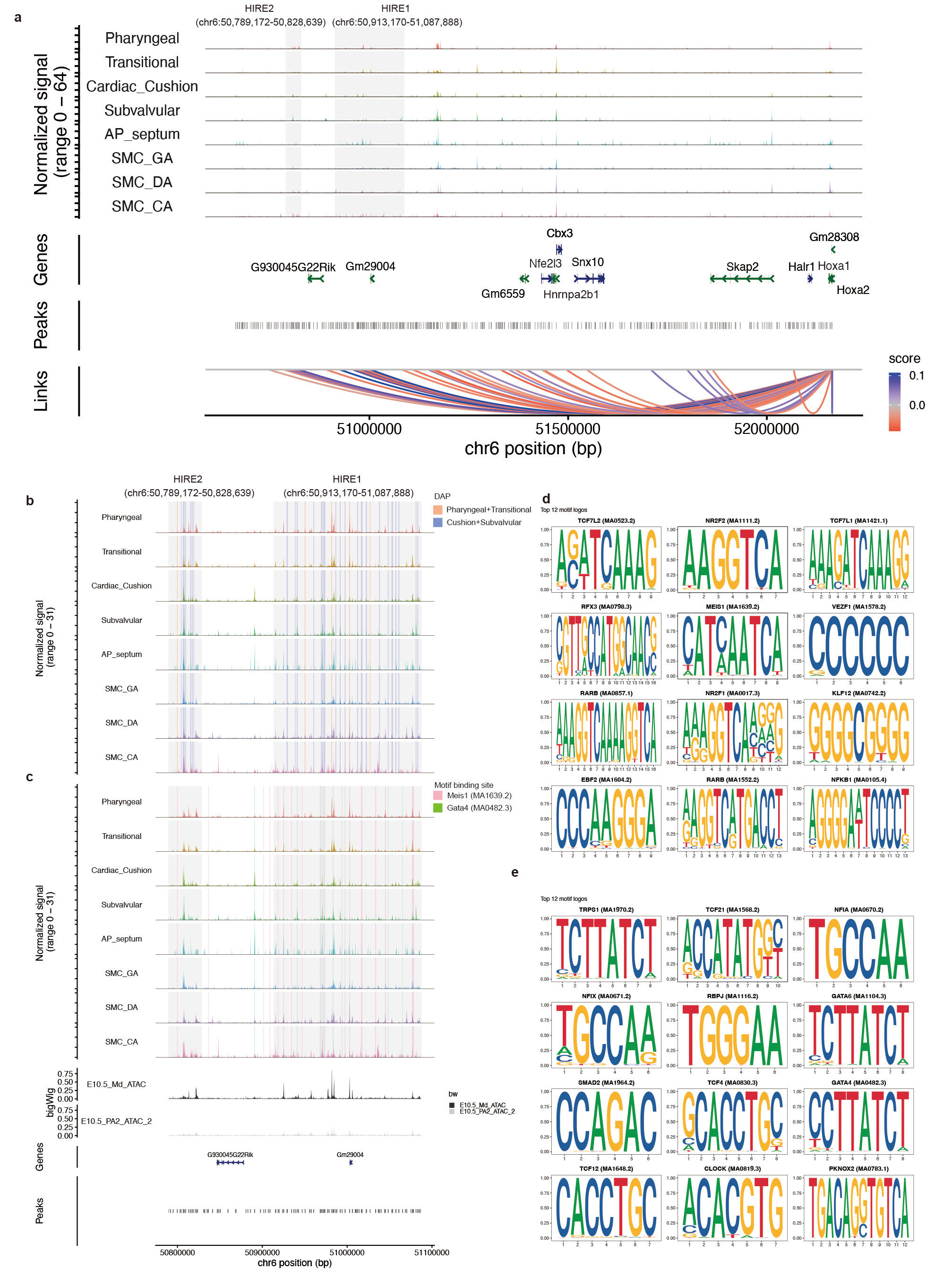

### Supplemental figure 12

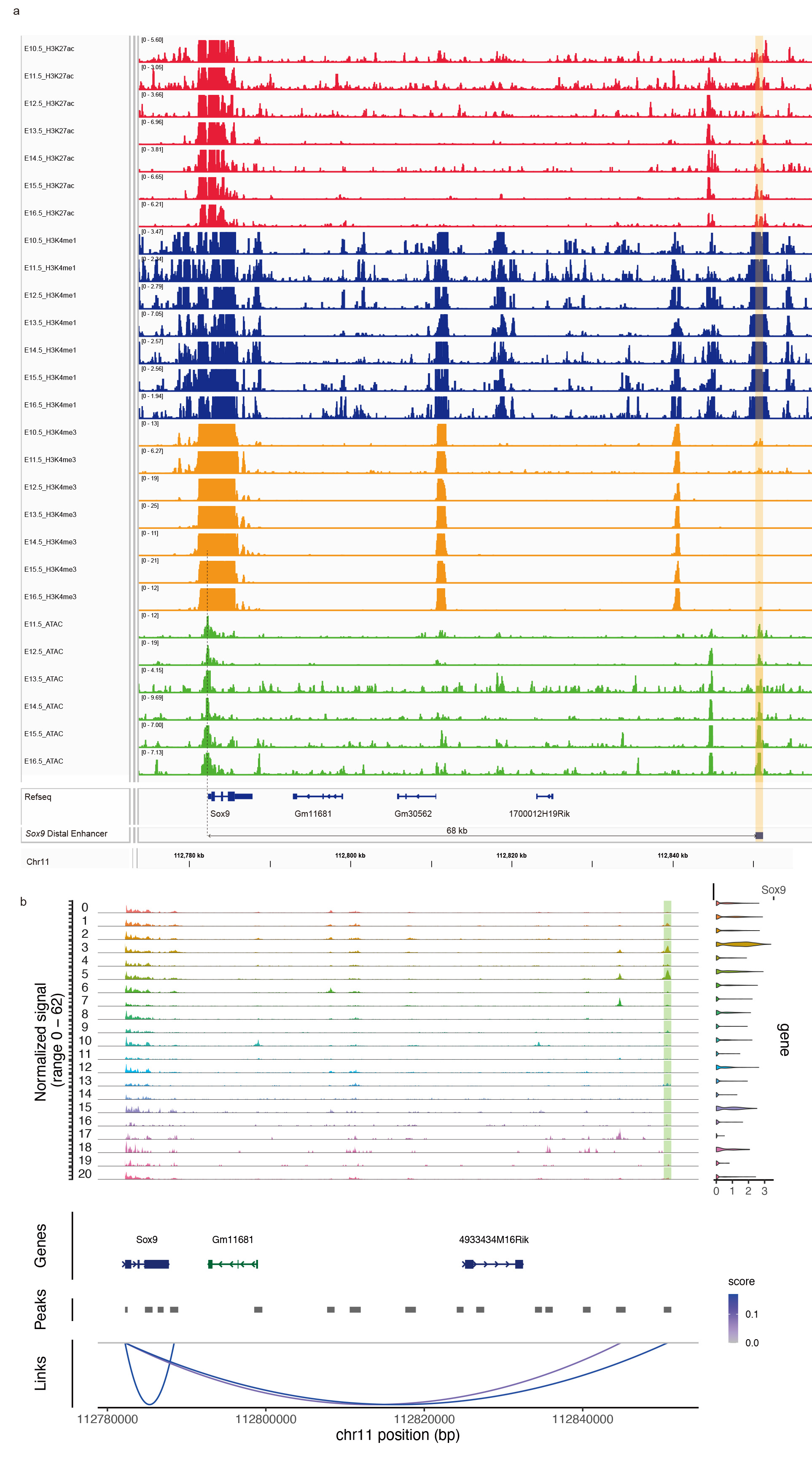

### Supplemental figure 13

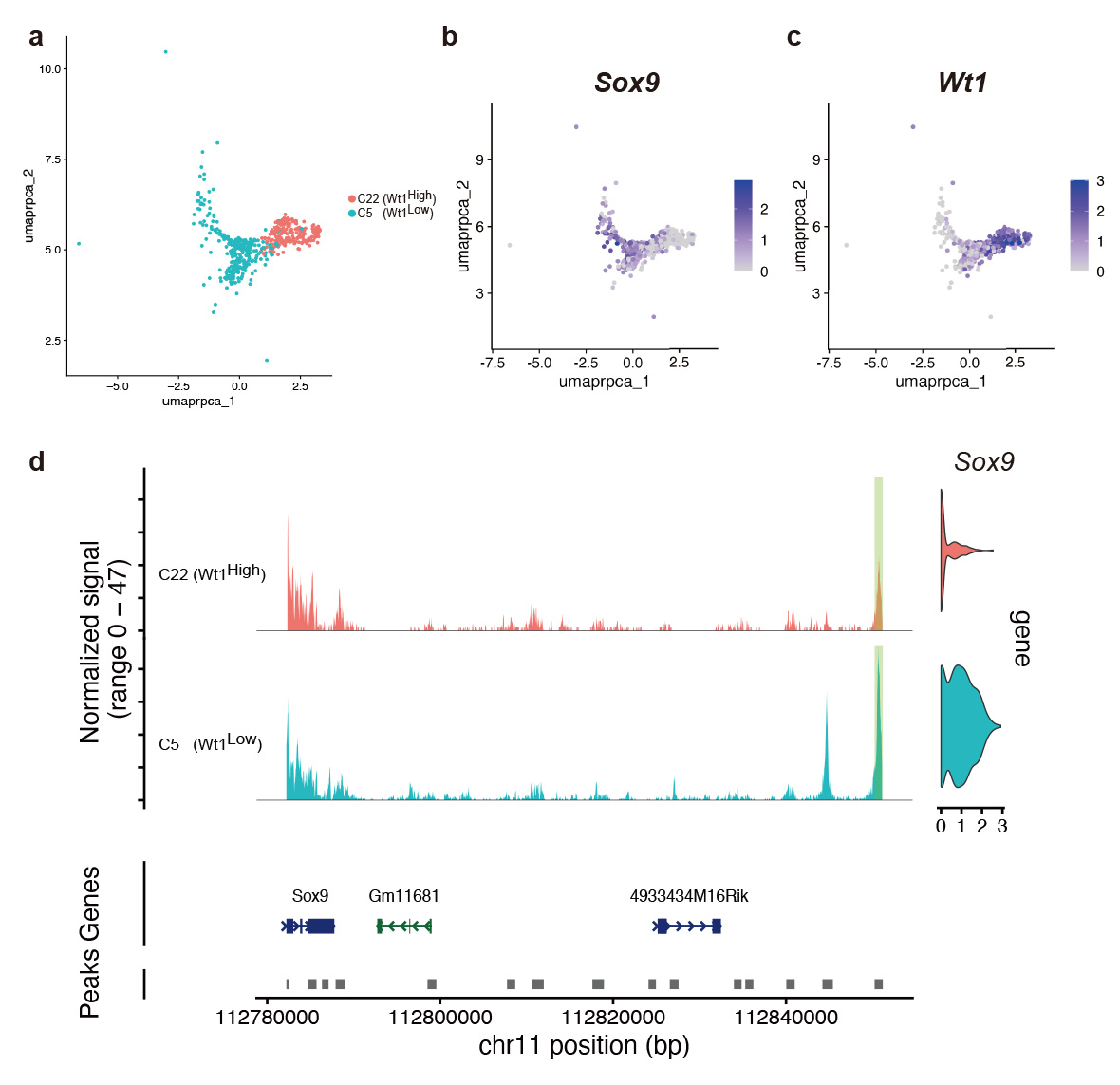
